# Whole-Genome Metagenomics Insights Revealed a Uranium Bio-remediating Cross-Domain Microbiome in the Dhala Impact Structure, India

**DOI:** 10.64898/2026.09.26.754616

**Authors:** Kapinder, Shivanshu Dwivedi, Anuj Kumar Singh, Jayanta Kumar Pati, Prateek Kumar

## Abstract

Meteorite impact structures on earth represent unique terrestrial analogues of extreme planetary environments where fractured lithologies, hydrothermal alteration, elevated concentrations of radionuclides, and prolonged water-rock interactions create ecological niches for specialized microbial communities. The Palaeoproterozoic Dhala impact structure in north-central India is characterized by uranium-bearing impactites and possesses a very high concentration of Uranium (up to 99.5 ppm) that makes it a unique natural laboratory to investigate microbial adaptation to radioactive and metal-rich geological environments. In addition, the Dhala structure harbors specialized radiotoxic hydrothermal mineral phases such as coffinite and pitchblende, which show a globally distinct and biologically unexplored niche. The present investigation is the first in-depth, whole-genome metagenomic study of soil and water samples of uranium-rich sites from the Dhala structure, exploring the whole microbial community (Bacteria, Archaea, Eukaryotes, and Viruses) up to the species level present in the samples. The cross-domain metagenomics analysis identified a highly unique and resilient microbiome that consists of bacteria, archaea, eukaryotes, and viruses, coevolved under prolonged radiological and multiple metal stress. Functional metagenomic analysis revealed numerous metabolic adaptations responsible for their survival, homeostasis, and biomineralization in situ. The deep sequencing method further enabled the identification of 58 functional genes directly involved in uranium bioremediation. Furthermore, this study revealed important pathways for radionuclide reduction, cellular efflux, and biotransformation. Novel findings of the present study point out the in-depth evolutionary relations of biosphere-geosphere interaction within an ancient hypervelocity impact-generated ecosystem. In a nutshell, the Dhala extremophiles with novel genetic makeup present an untapped and promising reservoir for the development of eco-friendly, next-generation microbial bioremediation strategies to efficiently tackle the global anthropogenic contamination of uranium and the challenges of nuclear waste management. Thus, the present study establishes Dhala structure as an unparalleled site for environmental geomicrobiology, microbial evolution, and astrobiology.

## 1. Introduction

Meteorite impacts have extensively influenced the geological, geochemical, and biological evolution of the Earth throughout its history [1,2,3]. Beyond the extreme terrestrial habitats generated by such catastrophic geological events, impact structures serve as typical sites for understanding geosphere-biosphere interactions and evolutionary limits of life [1,4,5,6,7,8]. They represent long-lived natural laboratories where hydrothermal alteration, fracturing and/or brecciation, and fluid-rock interactions collectively create highly heterogeneous subsurface environments capable of sustaining diverse microbial ecosystems [4,6,2,9,3]. These environments are characterized by steep physicochemical gradients, enhanced rock permeability, elevated mineral reactivity, and prolonged circulation of hydrothermal fluids, thereby generating ecological niches that are fundamentally different from those of surrounding crystalline basement rocks [10,2,3]. Therefore, terrestrial impact structures have increasingly attracted attention as exceptional geomicrobiological systems for investigating the interactions between geological processes and microbial life, while simultaneously serving as terrestrial analogues for potentially habitable environments on other planetary bodies, including Mars and the Moon.

Among naturally occurring extreme environments, the uranium-bearing geological systems occupy a particularly important position as they combine chemical toxicity, radioactivity, oxidative stress, and metal enrichment within a single ecological setting [11,12,13,14]. The naturally present radionuclide-rich ecological habitats put strong selection pressure on the indigenous microbial community [14]. Long-term exposure and stress from heavy metals and ionizing radiation drive the evolution of microbial communities at the genomic level. This results in the development of special metabolic pathways for biomineralization, metal tolerance, and radionuclide transformation. Identification of the genomic architecture of these extremophilic microbes is not only important for understanding their survival strategies, but it also provides information on novel genomic factors for the development of next-generation bioremediation methods for anthropogenic nuclear waste.

The Paleoproterozoic Dhala impact structure in India (25°17’51.42“N; 78°08’36.47”E), the largest impact crater in Southeast Asia, represents an exceptional natural analogue for such kinds of biogeochemical extremes (Fig. 1, Fig. 2) [15,16]. The Dhala structure was formed due to the hypervelocity impact of an ureilite-type meteorite on an Archean crystalline basement of the Bundelkhand craton [17,18,19]. Dhala is amongst the oldest and most deeply eroded meteorite impact structures on earth. This ∼25 km wide complex crater is characterized by a prominent central elevated area (CEA), extensive monomict breccia rings, and exposed outcrops of impact melt breccias [15,20,21,22]. Geochemical analyses in the literature have revealed that highly weathered descendant soils and local granitoids contain fractionated elevated base metals (Pb, Cu, Zn), distinct rare earth element (REE) patterns, and significant enrichment of hydrothermal ore minerals, including molybdenite, galena, and chalcopyrite [17,23].

**Figure 1:**
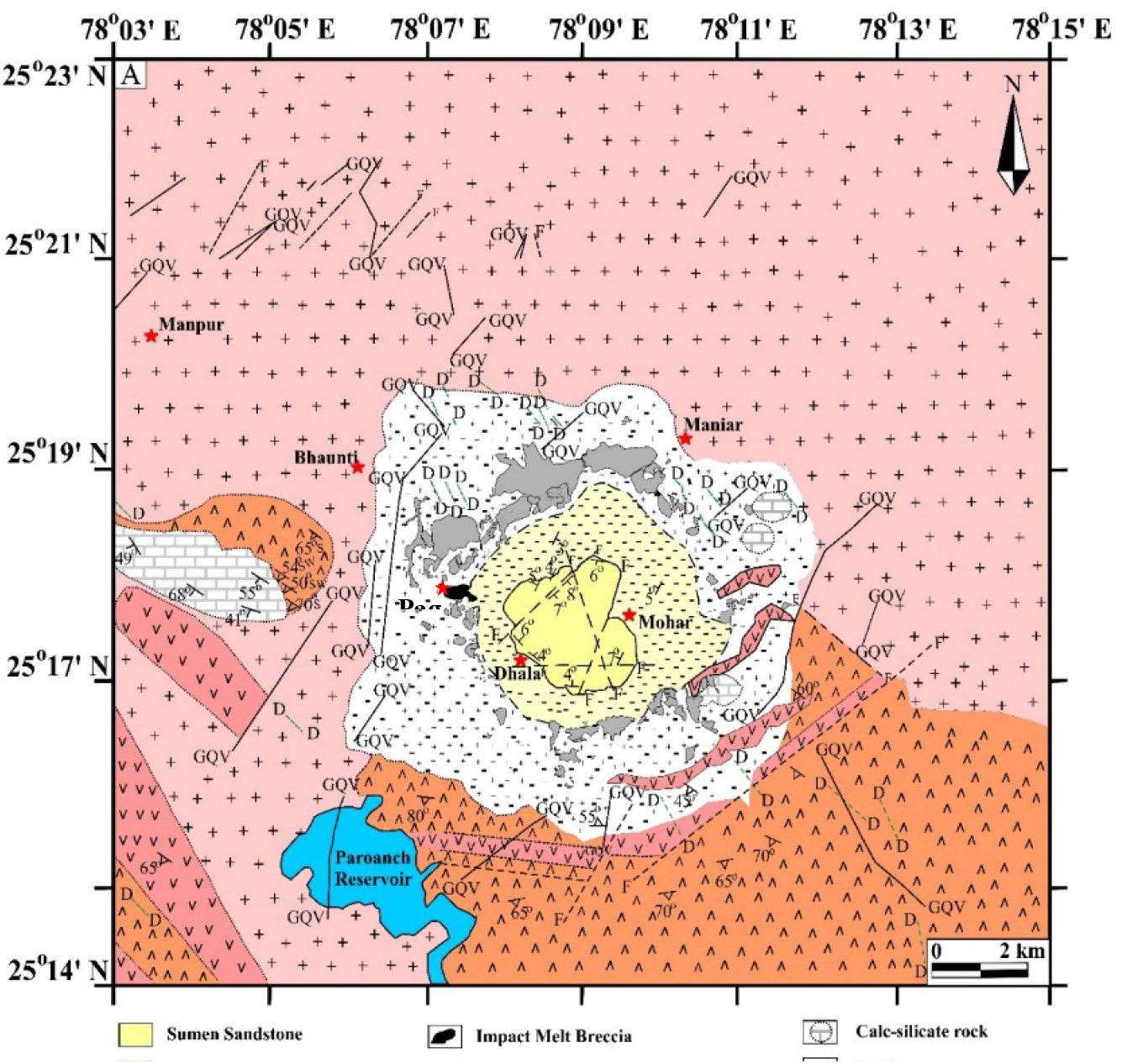
Geological map of the Dhala impact structure, India, showing the distribution of various rock types, settlements and water bodies present in the area (after Singh et al., 2021).

**Figure 2:**
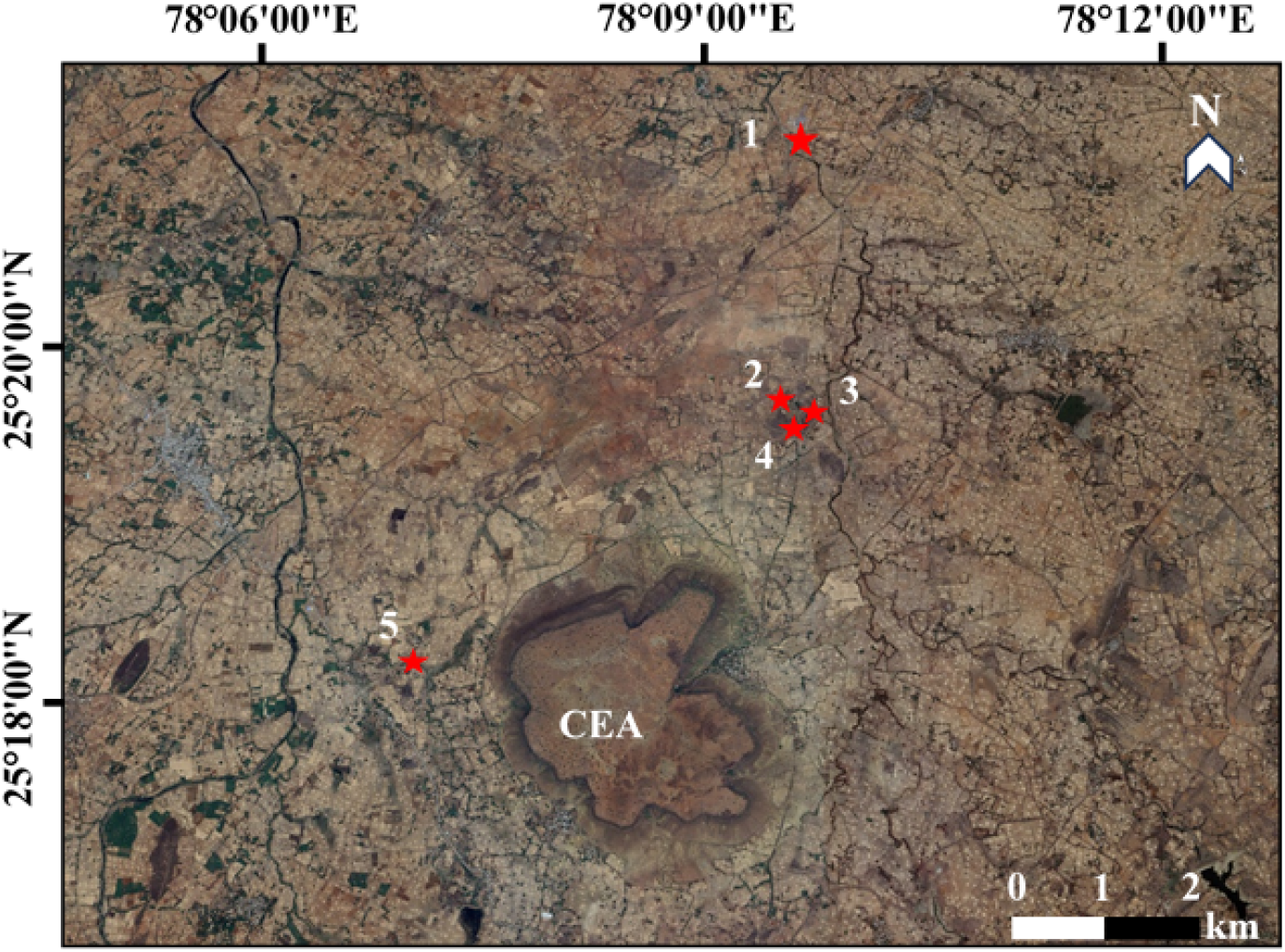
Google Earth image of the Dhala area, showing the Central Elevated Area (CEA) prominently. The 5^th^ red star denotes the location of the sampling site investigated in the present study. Image source: https://earth.google.com/web and date of acquisition: 08.09.2025.

Notably, the Dhala structure contains anomalous concentrations of thorium and uranium [24,25]. The uranium concentration reaches up to 99.5 ppm in pseudotachylite breccias and ranges from 8-40 ppm in the impact melt breccias [25]. Due to the metalliferous richness of the Dhala structure, the Atomic Minerals Directorate for Exploration and Research (AMDER), Government of India, conducted deep drilling campaigns for extensive subsurface geoscientific exploration of this site [15,26]. However, studies for the deep characterization of the indigenous biological landscape remain unexplored. An important knowledge gap in environmental geomicrobiology is represented by the biological ‘Dark Matter’ of the Dhala structure. The microbial population present in the water and soil habitats of this site has likely co-evolved under prolonged exposure to multiple metal and radiological stresses, resulting in the development of novel genomic architecture responsible for enhancing their survival and for bioremediating the heavy metals and radioactive compounds (e.g., uranium and thorium).

To uncover the complete microbial population (Bacteria, Archaea, Eukaryotes, and Viruses) of the Dhala structure and their adaptive mechanisms, the current study provides the first-ever report based on a comprehensive whole-genome metagenome. Using high-throughput sequencing, the present study extensively reveals the cross-domain biodiversity of this extreme habitat. The complex networks of bacteria, archaea, eukaryotes, and virus populations have also been explored. Further, functional genomic analysis provides information on metabolic factors performing heavy metal tolerance and in situ transformation. Most importantly, the present investigation identified 58 distinct genes directly associated with uranium bioremediation from the Dhala area. Findings of this study provide insights into microbial dynamics of an old, meteorite-impacted biosphere as well as identification of efficient genomic targets, which have direct application for eco-friendly bioremediation of global anthropogenic uranium contamination.

## 2. Materials and Methods

### 2.1 Study Location and Sampling Description

A field investigation of the Dhala structure, India, was conducted between 20^th^ and 24^th^ March, 2025. Sampling involved the collection of wet soil and water from the water body, selected based on their lithological characteristics and contamination levels of potential radioactive elements. From each site, two types of samples were collected separately: (i) ∼100 mL of water, stored in laboratory-grade plastic bottles, and (ii) wet soil samples, stored in airtight plastic bags.

The sampling site (25°18’9.09“N; 78°7’23.8”E) is located ∼2 km west of the CEA of the Dhala structure. It is an abandoned water well excavated within an outcrop of impact melt breccia (Fig. 3). The water in the well is dark green, stagnant, and polluted with decaying plant matter and anthropogenic debris, making it unfit for domestic or agricultural use. Both the soil and water in the well may exhibit elevated concentrations of radioactive contamination, consistent with previous reports on uranium- and potassium-bearing phases in the impact melt lithology of the Dhala structure. The water table at the site is relatively shallow. The surrounding soil is reddish brown, medium-to fine-grained, and derived predominantly from impact melt rocks. The collected boulders are vesiculated fragments of impact melt breccia.

**Figure 3:**
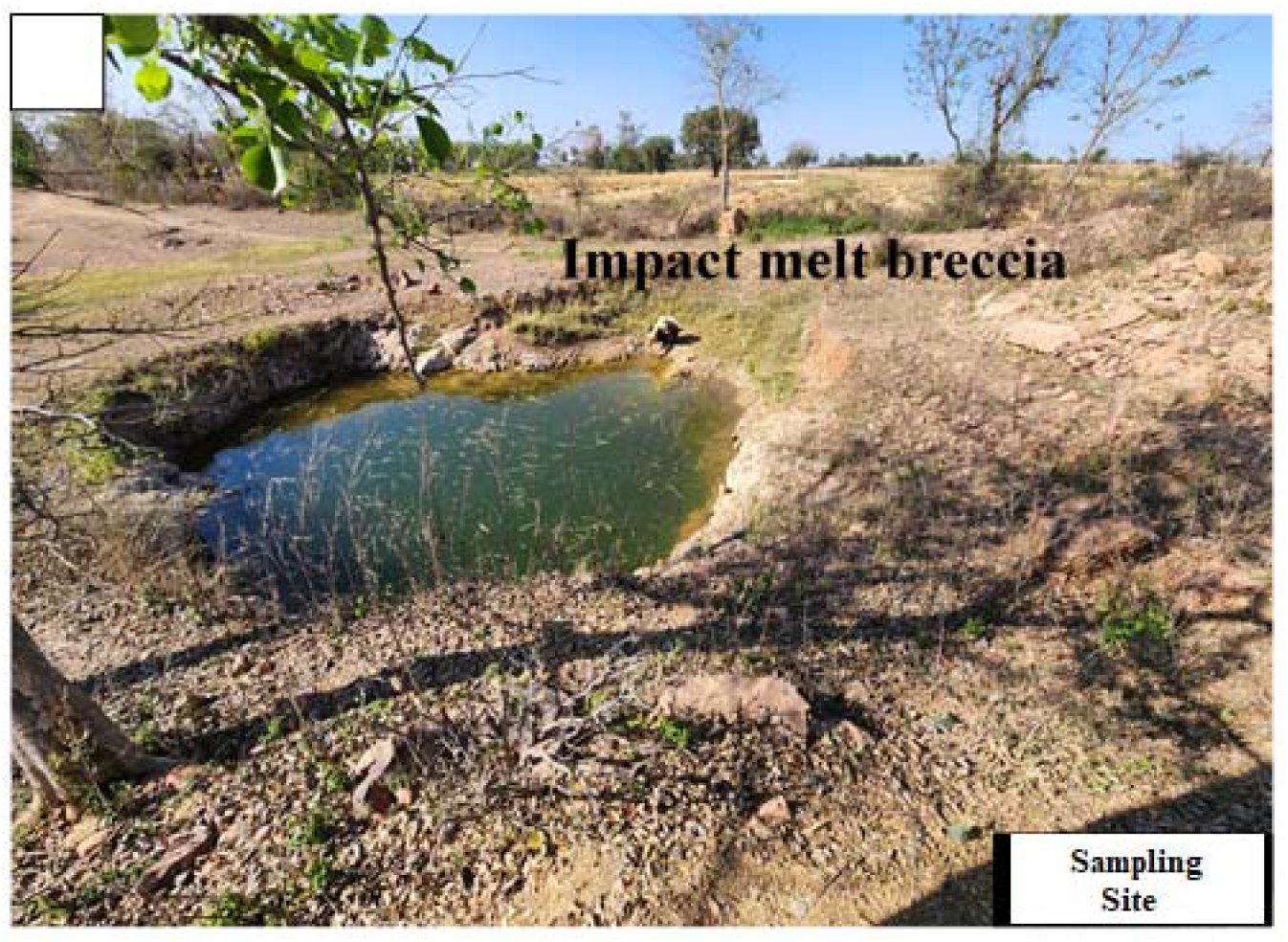
The field photograph of the sampling site within the Dhala impact structure, where soil and water samples were collected from/nearby the water body for the present study. This site is located about 2 km west of the CEA of the Dhala structure and shows a dug well developed entirely within the impact melt lithology.

### 2.2 Whole Metagenome Sequencing, Processing and Assembly of Raw Metagenomic Reads, Annotation, Taxonomic Analysis, Functional Analysis, and Data Visualization

Whole-genome metagenome sequencing of both soil (DS5) and water (DW5) samples from the Dhala area was performed using the Illumina NovaSeq-6000 platform. The quality of raw reads was assessed using FastQC v0.11.9 with default parameters [27]. Raw paired-end reads were processed using fastp v0.20.1 [28] with polyX trimming, sliding window quality filtering (4 bp), low-complexity read removal, and overlap-based base correction. Reads with an average Phred score below 10 or a length shorter than 60 bp were discarded. Additional trimming removed low-quality bases from the 5′ end (mean Q < 20) and the 3′ end (mean Q < 15). Processing was performed using 24 threads, and quality reports were generated in both HTML and JSON formats. A final quality reassessment was carried out using FastQC [27].

High-quality (HQ) reads were then annotated using Kraken2 v2.1.1 [29] against the RefSeq standard database (updated June 2024) (https://benlangmead.github.io/aws-indexes/k2). Taxonomic classification results were visualized using the Krona tool v2.8.1. The preprocessed HQ reads were assembled using metaSPAdes v3.15.5 [30] with k-mer sizes of 21, 33, and 55. Repeat resolution was enabled, while mismatch careful mode and Mismatch Corrector were disabled. Coverage cutoff was turned off to retain all contigs. The resulting assemblies were annotated using Prodigal [31] for gene prediction. Predicted genes were functionally annotated using EggNOG Mapper v2 [32] with the KEGG database, and the results were visualized using Krona v2.8.1.

The generated OTU table was rarefied to the minimum library size before comparative microbiome analysis. The relative abundance of bacterial, eukaryotic, and viral taxa was calculated from the phylum to species levels to identify dominant groups. The top 20 dominant genera and species across both groups were visualized using a heatmap. Alpha diversity metrics, including Chao1, Observed Species, Coverage, Pielou’s Evenness, Shannon, Simpson, Inverse Simpson, and Fisher’s Index, were calculated to assess species richness and evenness within microbial communities. A Venn diagram was constructed to identify common and unique taxa between groups. For functional analysis, the top 14 genes most abundant in uranium degradation were manually curated, sorted, and analyzed. All analyses were performed using the Microeco package in R [33].

## 3. Results

### 3.1 Identification of Bacterial and Archaeal Alpha Diversity

In alpha diversity results, the ACE (Abundance-based Coverage Estimator) index reveals that the soil sample of the studied site has more bacterial and archaeal species richness in contrast to the water sample collected from the Dhala structure (Fig. 4 (a)). Similarly, the results of Chao1 analysis demonstrated that the soil sample has greater species richness along with the presence of more rare species (Fig. 4 (b)). The elevated ACE and Chao1 values observed in the soil sample suggest the presence of a taxonomically complex microbial assemblage capable of exploiting a broader range of ecological niches. Such elevated richness may likely be associated with the heterogeneous mineralogical composition of the soil, which provides diverse microhabitats, redox gradients, and nutrient reservoirs capable of sustaining multiple microbial guilds simultaneously.

**Figure 4:**
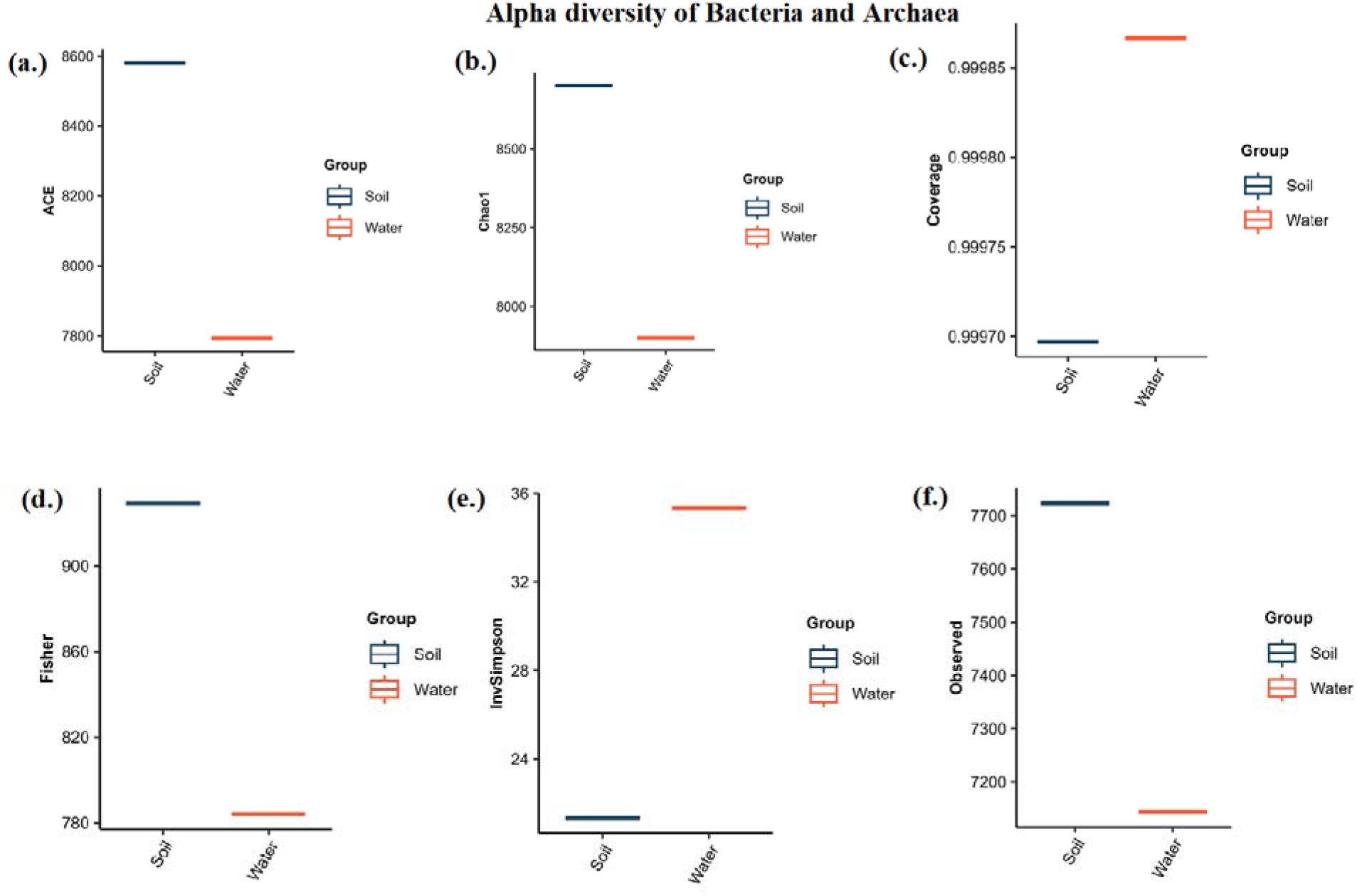
Alpha diversity analysis of Bacterial and Archaeal populations present in the soil and water samples collected from the Dhala area; **(a.)** ACE, **(b.)** Chao1 **(c.)** Coverage **(d.)** Fisher **(e.)** InvSimpson **(f.)** Observed.

The coverage graph explains how well or how completely the alpha diversity of microbes has been captured in each sample by sequencing (Fig. 4(c)). If the coverage is >0.95, it is a very good sampling, and estimation of alpha diversity is reliable. If the coverage is <0.90, then the sampling is not good, and alpha diversity could be biased in this case. In the present study, the coverage value exceeded 0.95 for both soil and water samples, indicating the correct estimation of alpha diversity, i.e., the sequencing depth was sufficient to capture the majority of the microbial diversity present in each habitat (Fig. 4 (c)).

The Fisher graph specifies microbial diversity in a given sample. In the present study, The soil sample exhibits relatively higher Fisher values than the water sample, indicating a more diverse bacterial and archaeal populations (Fig. 4 (d)). The elevated value obtained in the case of the soil sample further suggests higher species richness, more rare taxa, and the possibility of a more complex microbial community (Fig. 4 (d)).

In contrast, the Inverse Simpson (InvSimpson) index mainly helps in finding evenness and dominance of taxa in microbial communities, or in other words, it tells how balanced your microbial community is. Higher InvSimpson values are usually interpreted as evidence of presence of various bacteria and archaea, all are present in the same number, and no one is dominating. Therefore, it can be termed as a balanced and healthy community. In the InvSimpson graph of soil and water samples from the Dhala area, the later displays a higher value, indicating a more balanced and healthy community in the water sample (Fig. 4 (e)). Conversely, the comparatively lower InvSimpson value observed in the soil sample suggests an unequal/unbalanced amount of bacterial and archaeal population (Fig. 4 (e)). This indicate that although the soil environment supports a greater number of taxa overall, community abundance is concentrated within a subset of dominant microbial populations.

The observed graph expresses about different types of bacteria or archaea present in a sample The soil sample display a high bar value than the water sample, which means it contains more diverse array of bacteria or archaea (Fig. 4 (f)). This also indicates that there is a high richness and complex microbial community in the soil sample whereas, the low bar value of observed graph index for the water sample suggests presence of narrower range of microbes, low richness, and a disturbed environment (Fig. 4 (f)).

The community evenness analyses further demonstrated important ecological differences between the two habitats. Pielou’s evenness index plot represents how balanced or even the distribution of species is. In the present study, we constructed this plot to study the distribution pattern of bacteria and archaea in the Dhala samples (Fig. 5 (a)). In the soil sample, the value is <0.5, which means that there is dominance of a few taxa. In contrast, the higher evenness value (>0.5) observed in the water sample indicate that there is a moderate imbalance among bacterial and archaeal species (Fig. 5 (a)).

**Figure 5:**
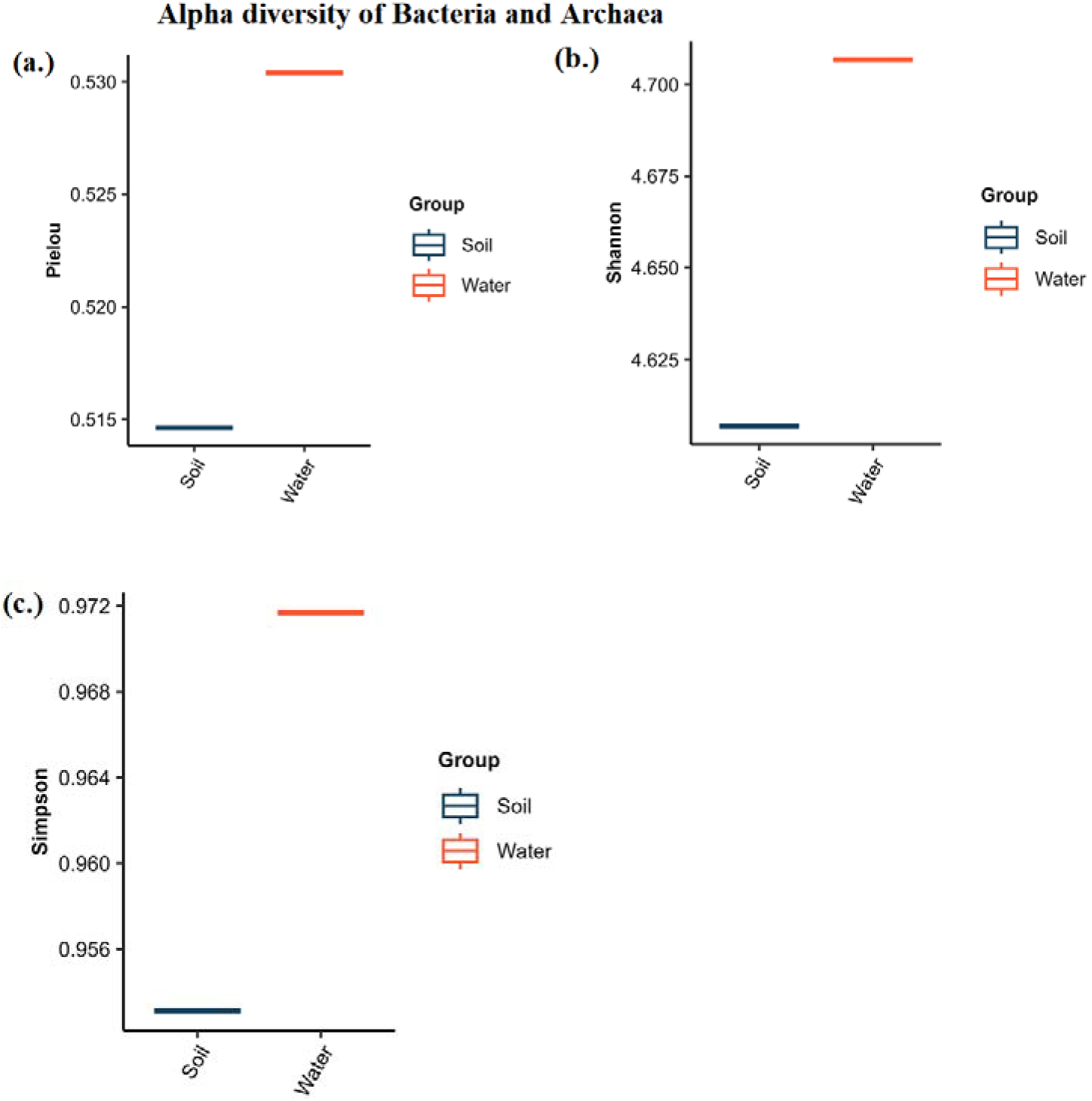
Alpha diversity analysis of Bacterial and Archaeal populations present in the soil and water samples collected from the Dhala area; **(a.)** Pielou, **(b.)** Shannon, **(c.)** Simpson.

The Shannon graph signifies the species diversity and richness. The soil sample exhibits moderate diversity of bacteria and archaea whereas the water sample characterizes a high diversity of bacteria and archaea (Fig. 5 (b)). In the Simpson graph, higher values indicate high diversity and lower values indicate low diversity. In the analyzed soil sample, the value was low, suggesting low diversity of bacteria and archaea. Conversely, a very high value for the water sample indicates a high diversity of bacteria and archaea (Fig. 5 (c)).

#### 3.1.1 Identification of Relative Abundance of Phylum, Class, Order, Family, Genus, and Species of Bacteria and Archaea in the Analyzed Soil and Water Samples of the Dhala Area

Across both the soil (DS5) and water (DW5) samples, bacterial communities exhibited a hierarchical taxonomic organization extending from the phylum to family level, with clear habitat-dependent differences in the relative abundance of individual taxonomic groups (Figs. 6-7). Metagenomic data analysis reveals that *Pseudomonadota* is the most abundant phylum (∼80%) in the soil sample, subsequently followed by phylum *Actinomycetota* (∼10%). Phylum *Myxococcota, Cyanobacteriota, Bacillota, Planctomycetota, Verrucomicrobiota, Acidobacteriota,* and others occupied the remaining fraction (10%) of the community. In the water sample, the phylum *Pseudomonadota* was most abundant (∼65%) followed by Phylum *Actinomycetota* (∼12%), and Cyanobacteriota (10%). Phylum *Bacillota, Euryarchaeota, Myxococcota,* and *Planctomycetota* occupy the rest proportion (13%) of the community (Fig. 6).

**Figure 6:**
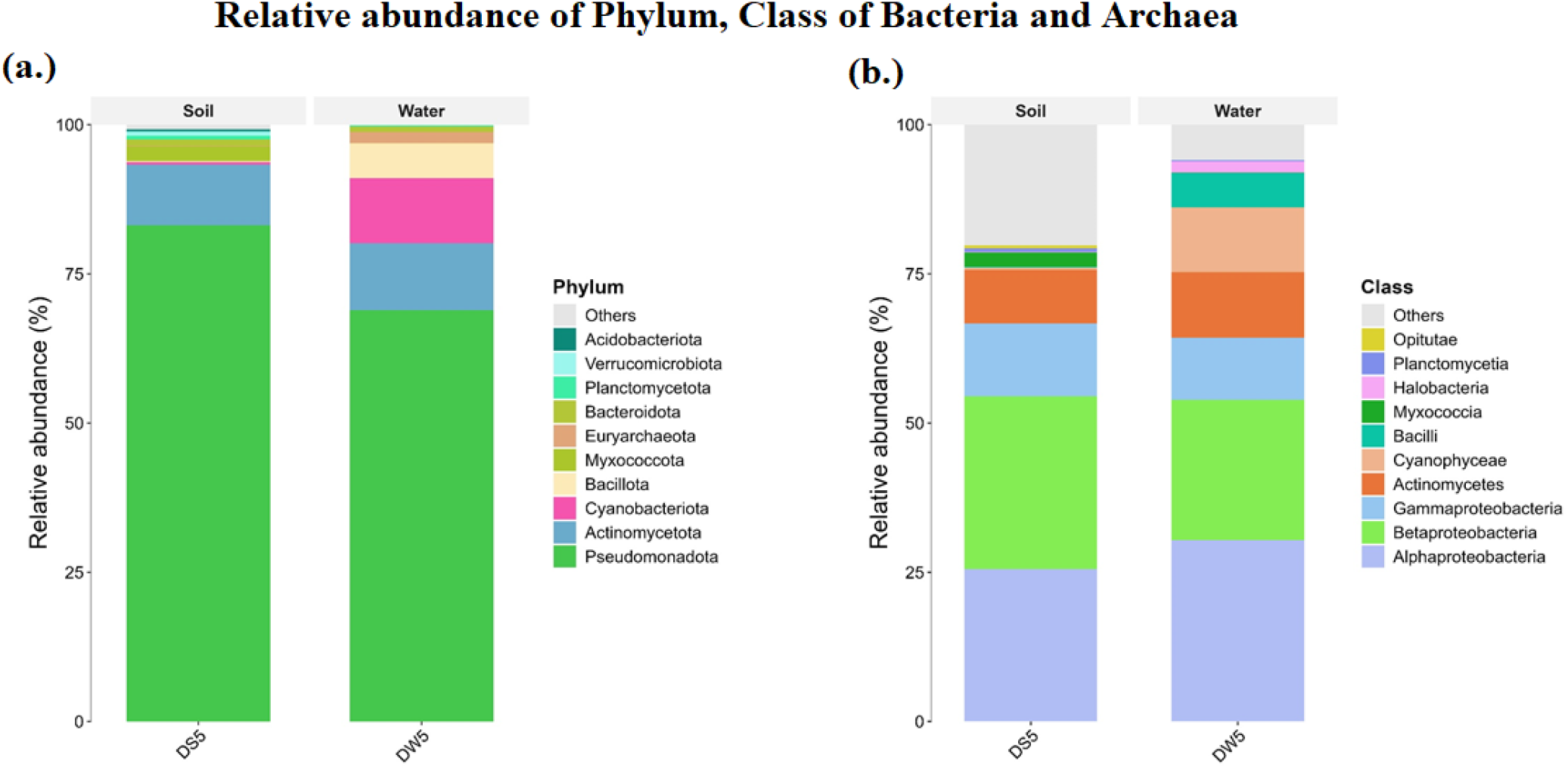
Stack bar plot showing the relative abundances of Phylum, Class of Bacteria, and Archaea in the soil and water metagenomes from the Dhala area.

**Figure 7:**
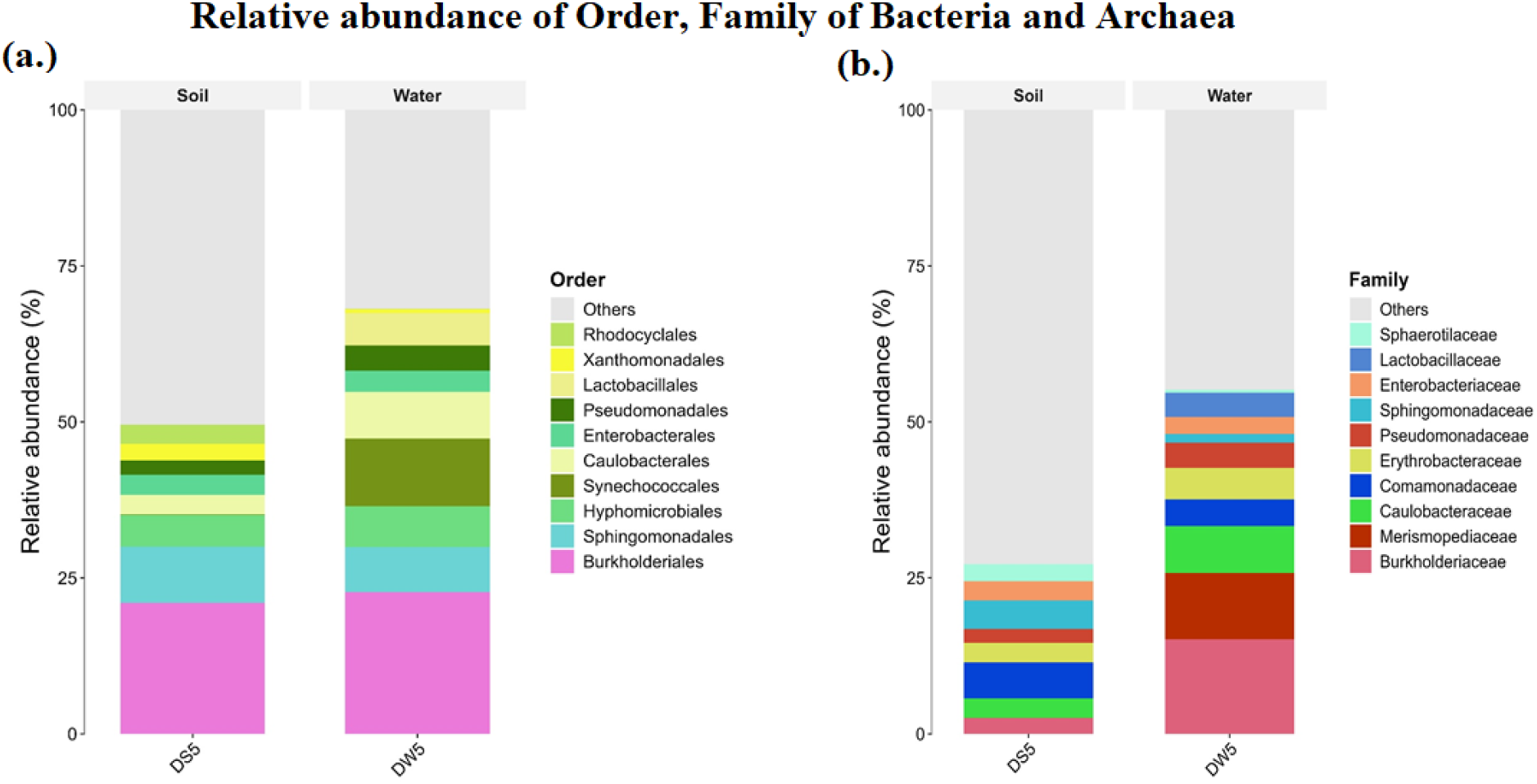
Stack bar plot depicting the relative abundances of Order, Family of Bacteria, and Archaea in the soil and water samples from the Dhala area.

The taxonomic resolution at the class level further highlighted the ecological organization of the bacterial community. The soil sample is dominated by *Betaproteobacteria, Alphaproteobacteria, Gammaproteobacteria,* and *Actinomycetes* whereas *Myxococcia, Cyanophyceae, Planctomycetia,* and *Opitutae* occur at relatively lower abundances (Fig. 6). A broadly similar taxonomic pattern was observed in the water sample, where *Alphaproteobacteria, Betaproteobacteria, Gammaproteobacteria, Actinomycetes,* and *Cyanophyceae* represent the principal classes while *Bacilli, Halobacteria,* and *Planctomycetia* are comparatively less abundant (Fig. 6). The predominance of the three major proteobacterial classes in both samples suggests that members of the *Pseudomonadota* constitute the principal taxonomic framework of the Dhala microbiome and likely perform many of the key ecological functions associated with elemental cycling, environmental adaptation, and radionuclide transformation.

The orders identified in the soil sample primarily include *Burkholderiales*, *Sphingomonadales, Hyphomicrobiales, Caulobacteriales, Enterobacterales, Pseudomonadales, Xanthomonadales, and Rhodocyclales* whereas in the water sample, major identified orders were *Burkholderiales*, *Synechococales, Hyphomicrobiales, Sphingomonadales, Caulobacteriales, Enterobacteriales, Pseudomonadales, and Lactobacillales* (Fig. 7).

During taxonomic analysis of the family in the soil sample, *Sphingomonadaceae, Comamonadaceae, Caulobacteraceae, Erythrobacteraceae, Merismopediaceae, Enterbacteriaceae,* and *Sphaerotilaceae* were identified. In the water sample, *Burkholderiaceae, Merismopediaceae, Caulobacteraceae, Comamonadaceae, Erythrobacteraceae, Pseudomonadaceae, Lactobacillaceae, Enterobacteriaceae, Sphingomonadaceae*, and *Sphaerotilaceae* families were primarily identified, and the rest were others (Fig. 7).

The genera identified in the soil sample includes *Synechocystis, Pseudomonas, Polynucleobacter, Polyphyrobacter, Sphingomonas, Bradyrhizobium, Caulobacter,* and the rest were others. In the water sample, genus *Synechocystis*, *Burkholderia, Polynucleobacter, Caulobacter, Pseudomonas, Lactobacillus, Porphyrobacter, Bradyrhizobium, Brevundimonas,* and *Sphingomonas* were commonly recognized, and the rest were others (Fig. 8).

**Figure 8:**
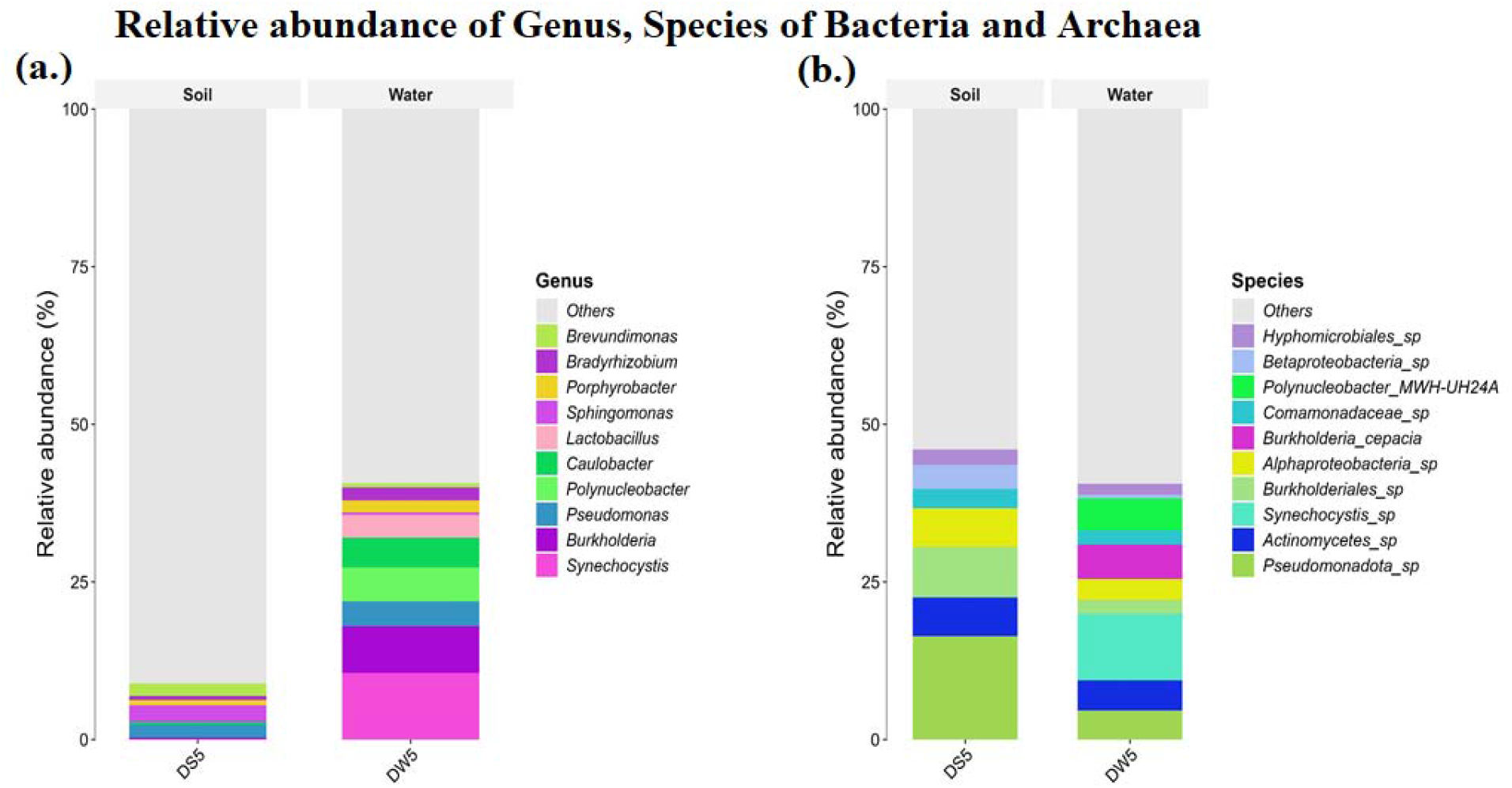
Stack bar plot displaying the relative abundances of Genus, Species of Bacteria, and Archaea in the soil and water metagenomes from the Dhala area.

Species of diverse genera such as *Pseudomonadota* sp., *Burkholderiales* sp., *Actinomycetes* sp., *Alphaproteobacteria* sp., *Betaproteobacteria* sp., *Synechocystis* sp., *Hypomicrobiales* sp. were identified in the soil sample,, and the rest were others. Whereas in the water sample, species of genus *Synechocystis* sp., *Polynucleobacter* MWH-UH24A, *Burkholderia cepacia, Pseudomonadota* sp., *Alphaproteobacteria* sp., *Comamonadaceae* sp., *Burkholderiales* sp., *Hypomicrobiales* sp., *Betaproteobacteria* sp. were identified, and the rest were others (Fig. 8).

#### 3.1.2 Identified dominant genus and species of Bacteria and Archaea in soil (DS5) and water (DW5) samples, with several having Uranium remediation potential

Although both the soil (DS5) and water (DW5) samples from the Dhala area share a common taxonomic framework, marked differences in the relative abundance of individual genera reflect habitat-specific ecological selection within the (uranium-bearing target) impact environment. Based on whole-genome metagenomics data, heat maps for the top 20 most abundant genera and species of Bacteria and Archaea in the soil and water samples were constructed (Fig. 9). Amongst the identified taxa, Genus *Pseudomonas* is dominant in both soil and water (DS5 and DW5) samples whereas the broader group *Pseudomonadota* sp. exhibits high abundance in the soil sample. *Pseudomonas* is well known for its heavy metal bioremediation ability, rapid physiological adaptation, efficient biofilm formation, and to actively precipitate Uranium (i.e., biomineralization) (Fig. 9).

**Figure 9:**
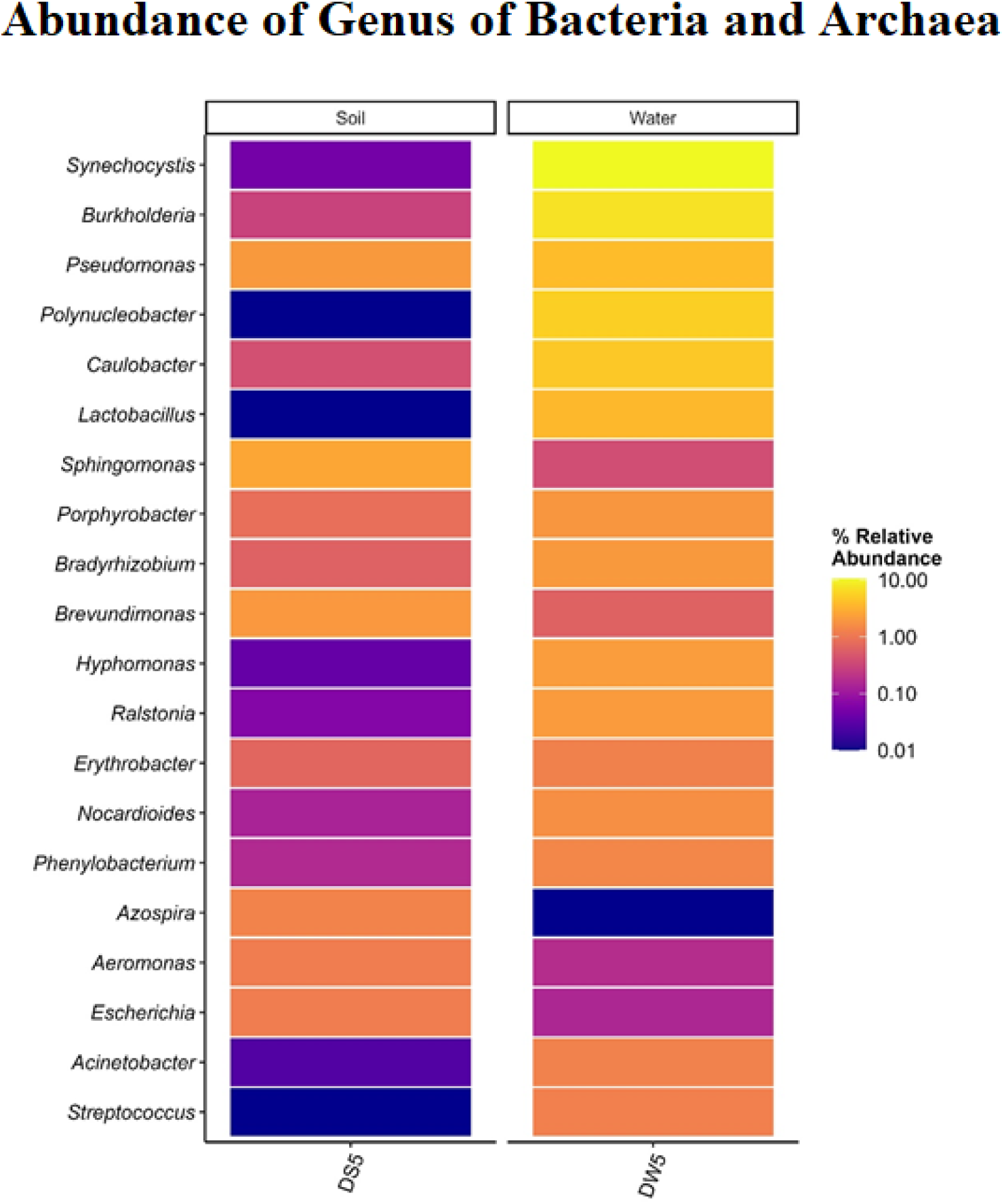
Heat map exhibiting the distribution of the top 20 Genus of Bacteria and Archaea identified in soil (DS5) and water (DW5) samples collected from the Dhala area. Each row in the heat map represents a different genus.

On the other side, Genus *Synechocystis* and *Synechocystis* sp. were abundant in the water sample (DW5). This genus and species of this genus are well-adapted photosynthetic cyanobacteria. They effectively perform biosorption, in which the heavy metals such as Uranium from aquatic habitats are accumulated and trapped directly on the cell walls (Figs. 9-10).

**Figure 10:**
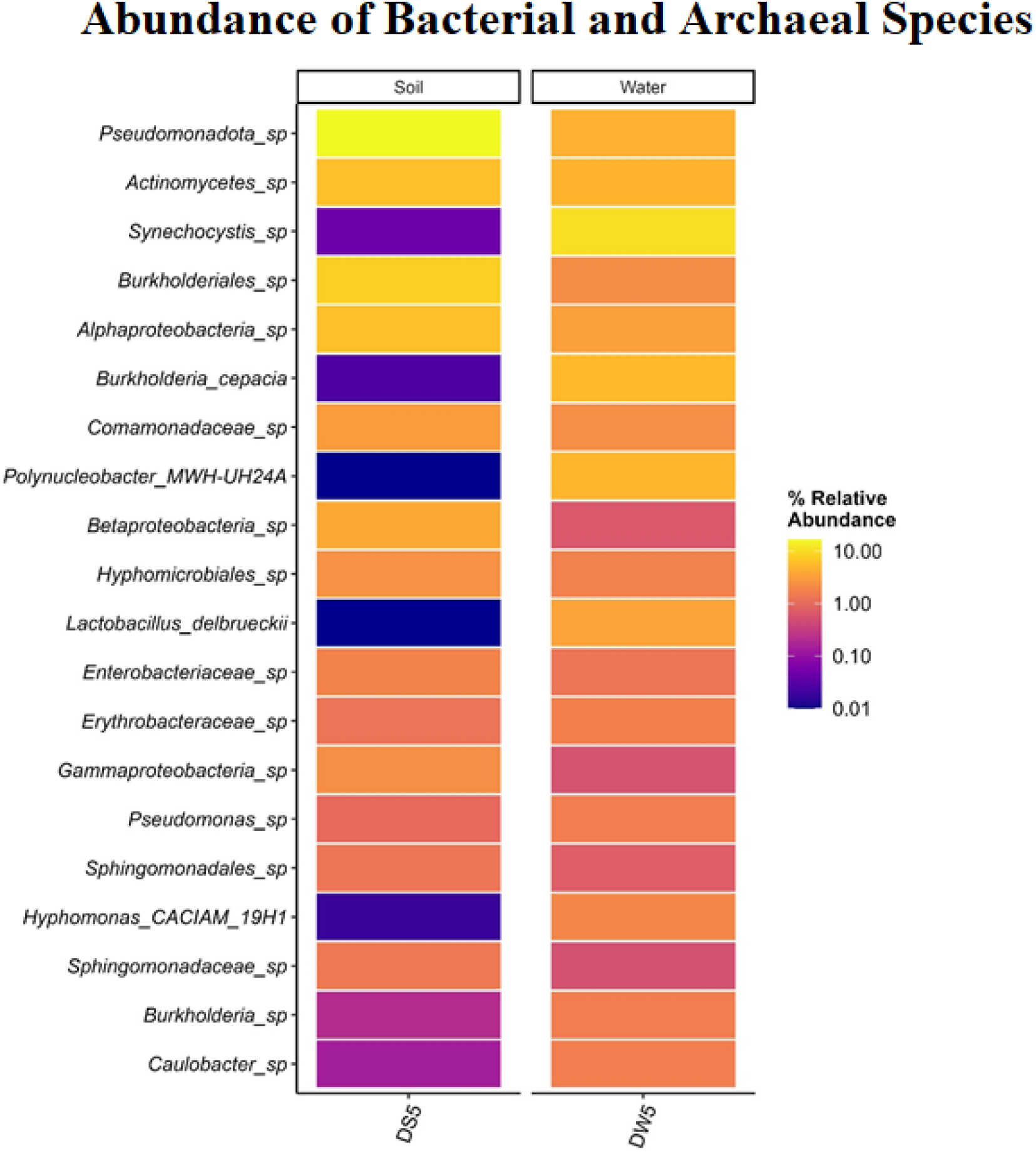
Heat map showing the distribution of top 20 Species of Bacteria and Archaea identified in soil (DS5) and water (DW5) samples from the Dhala area. Each row in the heat map represents a different Species.

Genus *Burkholderia* and specific species *Burkholderia cepacia* were more enriched in the water sample (DW5) in comparison to the soil sample (DS5). They possess a metal-efflux pump, which help in tapping of soluble Uranium (Figs. 9-10). Their persistence in both terrestrial and aquatic habitats indicates successful adaptation to the long-term physicochemical conditions generated by impact-induced fracturing, hydrothermal alteration, and uranium mineralization.

Genus *Sphingomonas* was detected in moderate amounts in both soil and water samples. It possesses a highly impermeable and unique outer cell membrane. Microbes from this genus are utilized in the bioremediation of Uranium through binding and biomineralizing it at the cell surface (Fig. 9).

### 3.2 Identification of Eukaryotic Alpha Diversity

Alpha diversity analyses revealed clear differences in the composition and diversity of eukaryotic communities between the soil and water samples collected from the Dhala impact structure (Figs. 11-12). The ACE (Abundance-based Coverage Estimator) index revealed that the water sample of the studied site has more eukaryotic species richness than the soil sample (Fig. 11(a)). The result of Chao1 analysis, which indicates the total number of species in a sample and is helpful in the estimation of rare/unobserved species in an analyzed sample, further revealed that the water sample has more eukaryotic species richness alongwith the a greater proportion of more rare species. The coverage graph explains how well or how completely the alpha diversity of microbes has been captured in each sample by sequencing. If the coverage is >0.95, it is a very good sampling, and estimation of alpha diversity is reliable. If the coverage is <0.90, then the sampling is not good, and alpha diversity could be biased in this case (Fig. 11(b)). In the present study, the coverage value exceeds 0.95 for both the soil and water samples, indicating the correct estimation of alpha diversity (Fig. 11(c)). In contrast to the richness estimators, Fisher graph (indicates microbial diversity in a given sample) showed that the soil sample contains a very high diversity of eukaryotic populations than the water sample, suggesting high species richness, more rare taxa, and the possibility of a more complex eukaryotic microbial community (Fig. 11(d)).

**Figure 11:**
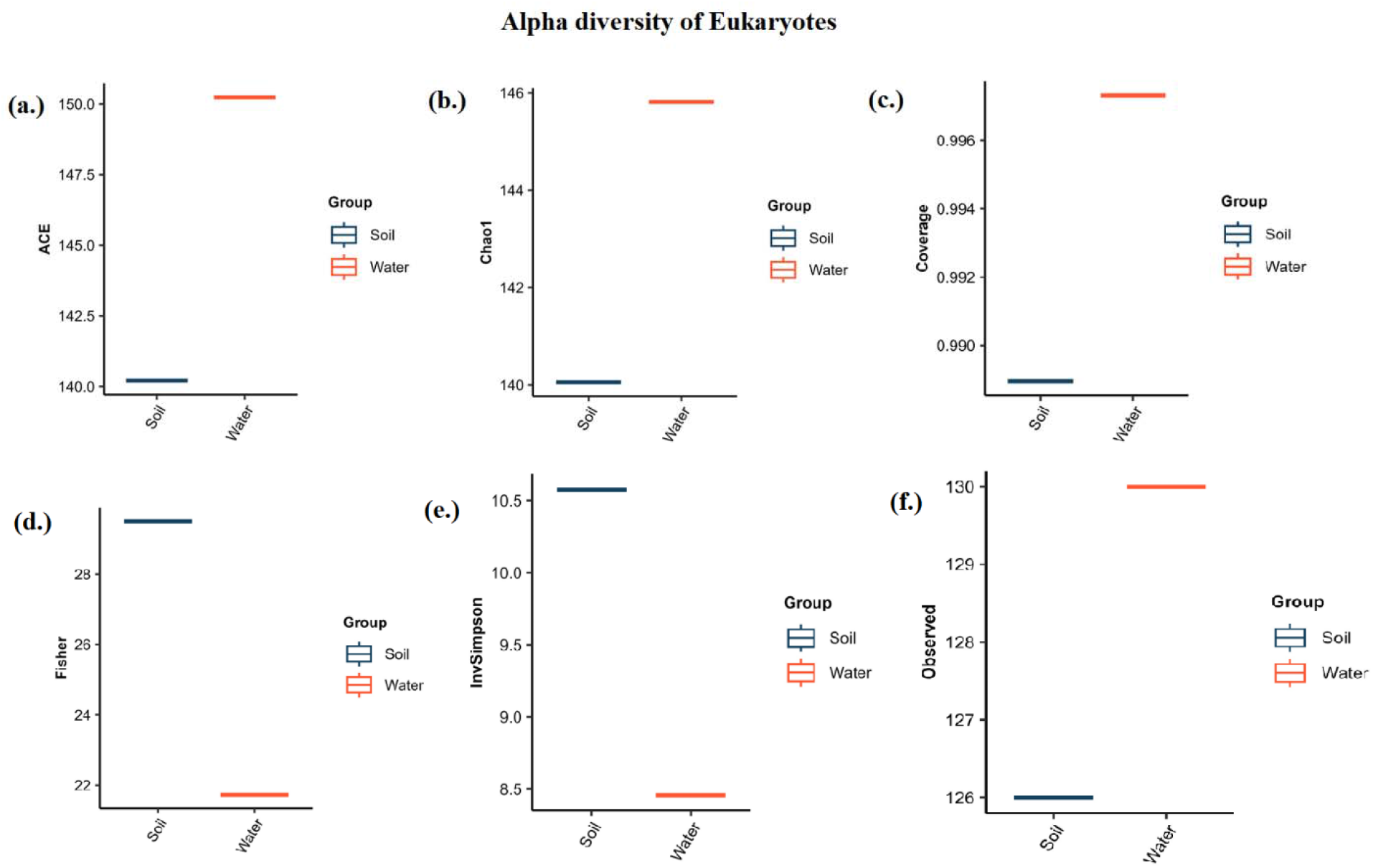
Alpha diversity analysis of Eukaryotic population present in the soil and water samples collected from western part of the Dhala structure; **(a.)** ACE, **(b.)** Chao1 **(c.)** Coverage **(d.)** Fisher **(e.)** InvSimpson **(f.)** Observed.

**Figure 12:**
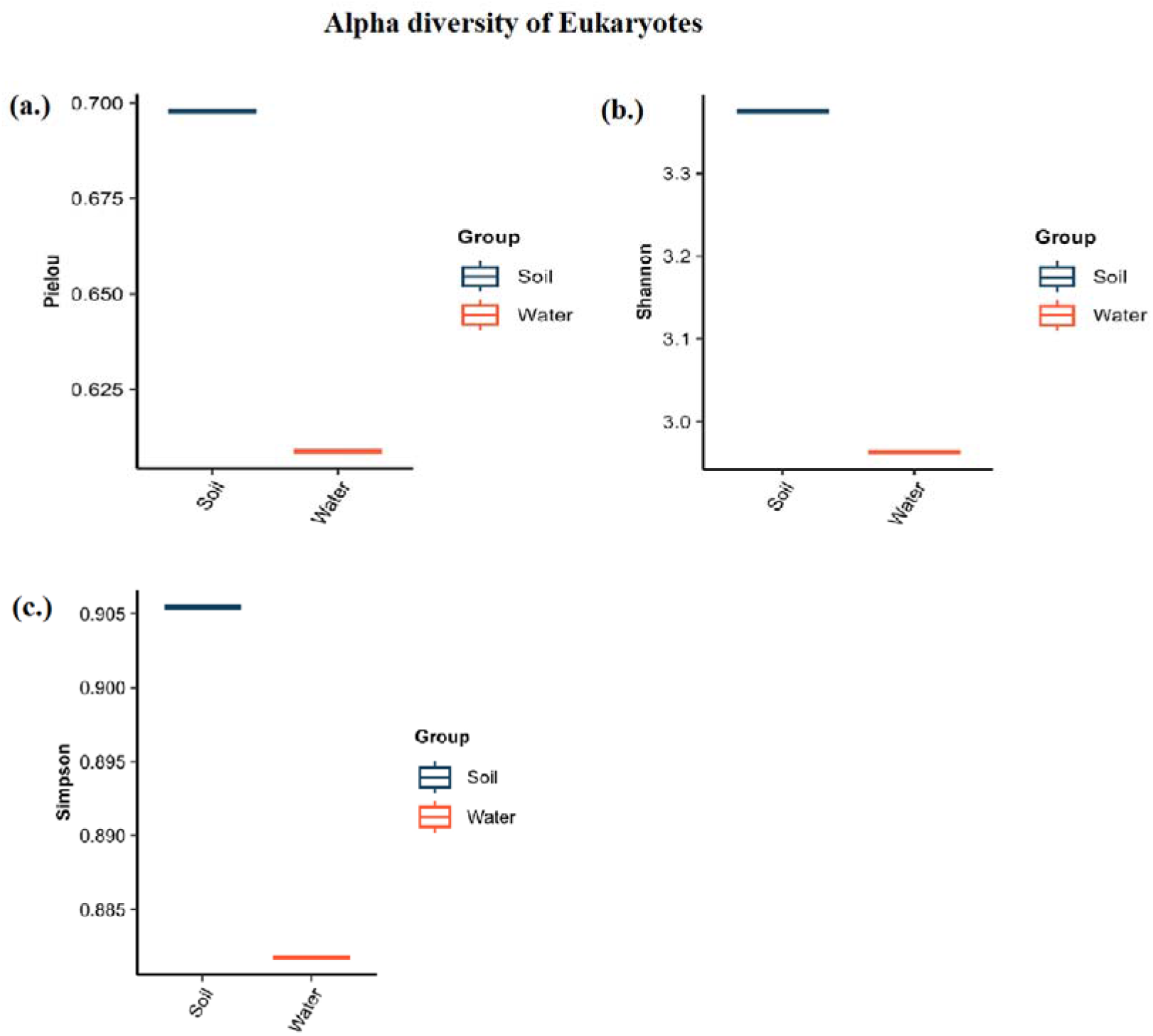
Alpha diversity analysis of the Eukaryotic population present in the soil and water sample collected from the Dhala area; **(a.)** Pielou, **(b.)** Shannon, **(c.)** Simpson.

The Inverse Simpson (InvSimpson) graph mainly helps in finding evenness and dominance of taxa in microbial communities, or in other words, it tells how balanced your microbial community is. If there is a high InvSimpson value, that means there is a presence of various eukaryotes, all are present in the same number, and no one is dominating. Therefore, it can be termed as a balanced and healthy community. In the present study, the InvSimpson graph of eukaryotes shows a higher value for the soil sample, reflecting a more balanced and healthy community. On the other side, a very low value for the water sample in the InvSimpson graph suggests an unequal/unbalanced amount of eukaryotic population present in the sample (Fig. 11(e)). The observed graph tells about how many different types of eukaryotes are present in any sample. The high bar value for the water sample means that it contains a more rich and complex eukaryotic microbial community than the soil sample (Fig. 11(f)).

Pielou’s evenness index plot represents how balanced or even the distribution of species is. In the present study, we constructed this plot to study the distribution pattern of eukaryotes. The soil sample a higher evenness value (0.700), which indicates a uniform distribution of eukaryotic species, whereas the water sample displayed a moderately lower value (>0.600), suggesting a less even eukaryotic species distribution (Fig. 12(a)). The Shannon graph, which explicits the species diversity and richness, further highlighted these compositional differences by recording higher diversity of eukaryotes in the soil sample than in the water sample (Fig. 12(b)). Similarly, the Simpson diversity graph showed higher values for the soil sample, confirming that the terrestrial habitat supports a more diverse and evenly distributed eukaryotic community, whereas the aquatic habitat is characterized by lower diversity and greater taxonomic dominance (Fig. 12(c)).

#### 3.2.1 Identification of Relative Abundance of Phylum, Class, Order, Family, Genus, and Species of Eukaryotes in the Analyzed Soil and Water Samples

The phyla *Ascomycota, Bacidiomycota, Chordata,* and *Euglenozoa* are abundant in the soil sample with minor occurrence of *Bacillariophyta*. In the water sample, the common phyla identified include *Ascomycota, Basidomycota,* and *Bacillariophyta* while *Fornicate, Apicomplexa, Euglenozoa, Evosea,* and *Cerozoa* are less abundant. Major classes documented in the soil sample are *Dothideomycetes, Sordariomycetes, Mammalia, and Kinetoplastea* followed by small proportion of *Eurotiomycetes, Saccharomycetes, Aconoidasida, Conoidasida, Malasseziomycetes, and Bacillariophyceae*. Conversely, *Malasseziomycetes, Saccharomycetes,* and *Eurotiomycetes* were abundant classes identified in the water sample, followed by minor occurrences of *Dothideomycetes, Mammalia, Kinetoplastea, Aconoidasida, Bacillariophyceae,* and *Conoidasida* classes (Fig. 13 (a,b)).

**Figure 13:**
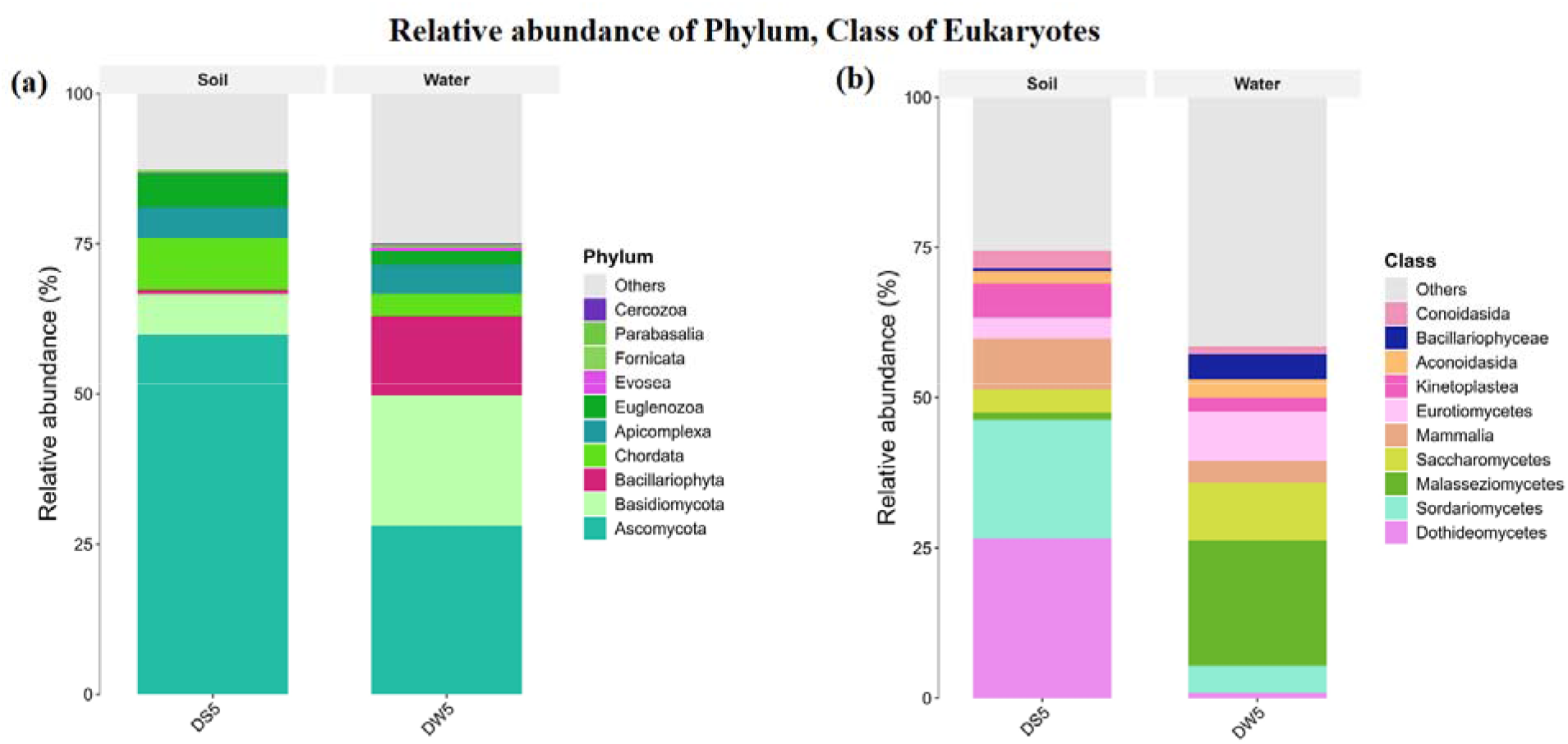
Stack bar plot showing the relative abundance of the Phylum, Class of Eukaryotes in the soil and water samples from the Dhala area.

In the soil sample, the dominant orders included *Mycosphaerellales, Primates, Hypocreales,* and *Trypanosomatida,* whereas *Saccharomycetales, Malasseziales, Eurotiales, Eucoccidiorida,* and *Thalassiosirales* were detected at comparatively lower abundance. In contrast, the water sample was characterized by the predominance of *Malasseziales, Saccharomycetales, Eurotiales,* and *Trypanosomatida,* while *Primates, Thalassiosirales, Hypocreales,* and *Eucoccidiorida* were relatively less abundant (Fig. 14(a,b)).

**Figure 14:**
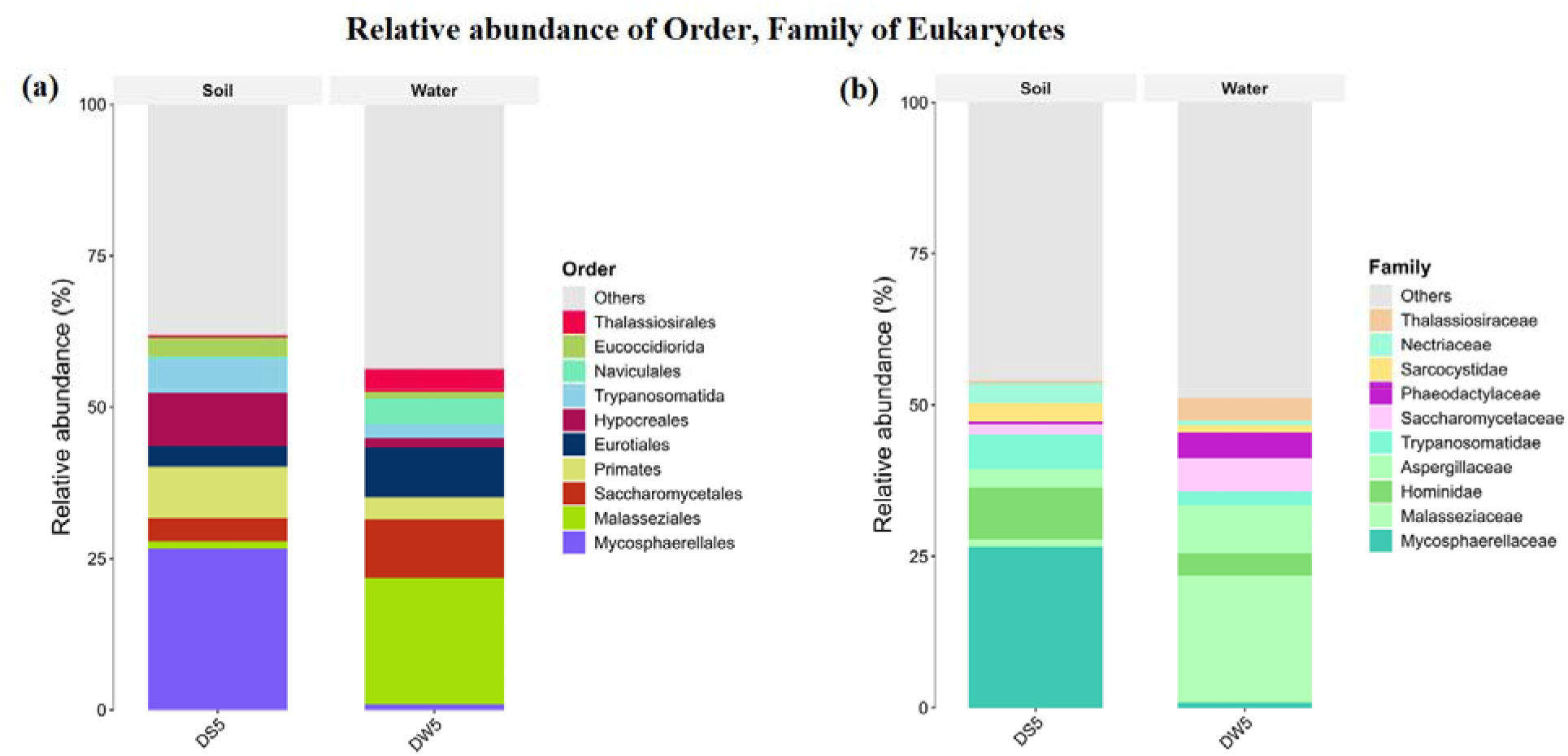
Stack bar plot showing the relative abundance of Order, Family of Eukaryotes

A similar pattern was observed at the family level. The soil sample was dominated by *Mycosphaerellaceae, Hominidae,* and *Trypanosomatidae*, whereas *Malasseziaceae, Aspergillaceae, Saccharomycetaceae, Phaeodactylaceae, Sarcocystidae, Nectriaceae,* and *Thalassiosiraceae* occurred at lower relative abundances. The water sample exhibited enrichment of *Malasseziaceae, Aspergillaceae,* and *Saccharomycetaceae* families, while *Hominidae, Trypanosomatidae, Phaeodactylaceae, Nectriaceae,* and *Thalassiosiraceae* were comparatively underrepresented (Fig. 14(a,b)).

The soil sample was characterized by abundance of *Cercospora, Homo, Leishmania, Fusarium,* and *Collectotrichum* genus, while *Malassezia* and *Aspergillus* were relatively less abundant. However, the water sample exhibited enrichment of *Malassezia, Aspergillus,* and *Phaeodactylum* were abundant, while *Homo, Leishmania, Fusarium,* and *Babesia* were comparatively underrepresented (Fig. 15(a,b)).

**Figure 15:**
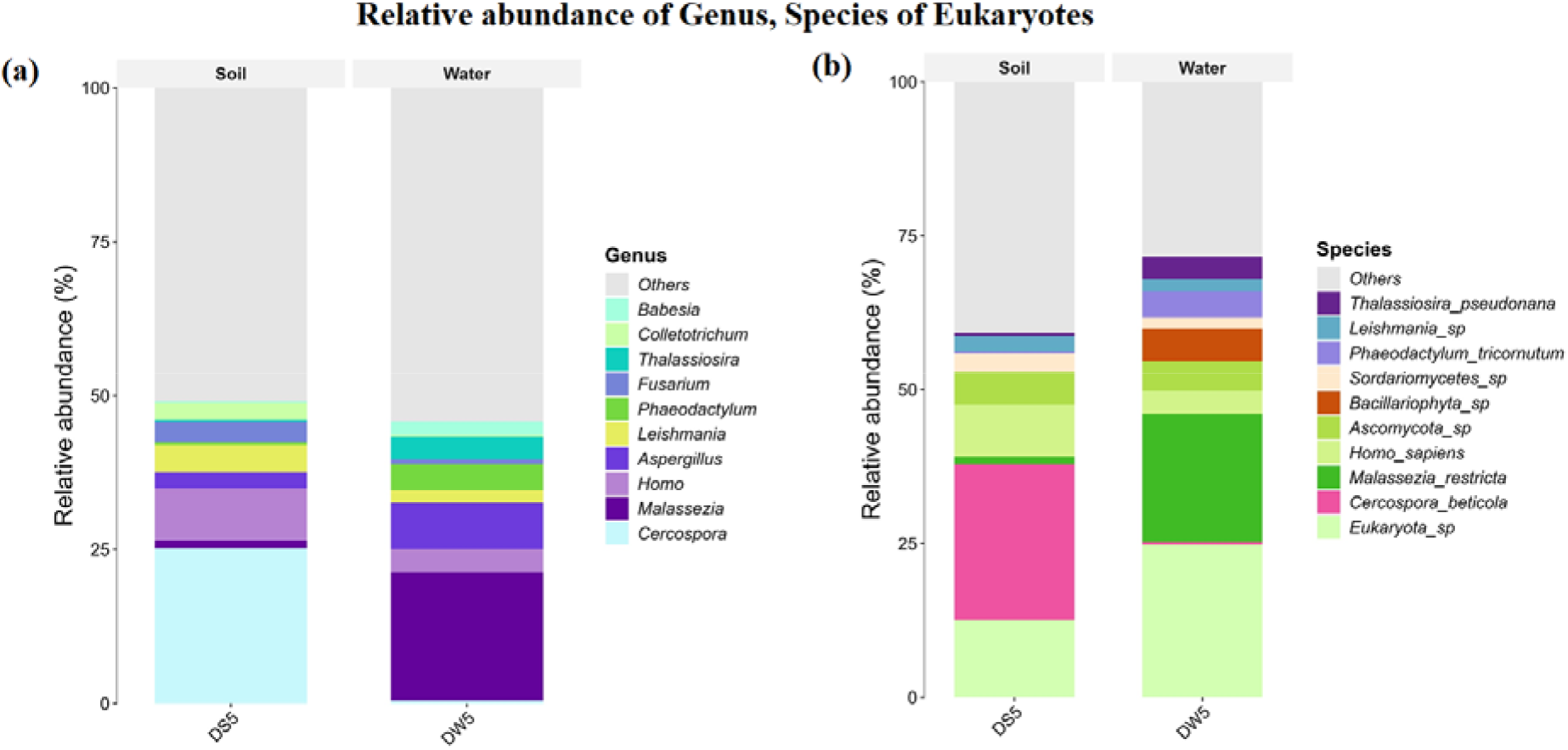
Stack bar plot showing relative abundance of Genus, Species of Eukaryotes

During species identification, the soil sample was dominated by *Cercospora beticola, Eukaryota* sp., and *Ascomycota* sp. with relatively lower abundances of *Malassezia restricta, Sordariomycetes* sp., *Leishmania* sp., and *Thalassiosira pseudonana*. The water sample exhibited a distinct species composition, with *Eukaryota* sp., *Malassezia restricta, Ascomycota* sp., *Bacillariophyta* sp., *Phaeodactylum tricornutum,* and *Thalassiosira pseudonana* representing the most abundant taxa. However, *Cercospora beticola, Sordariomycetes* sp., and *Leishmania* sp. occurred at relatively low abundances (Fig. 15(a,b)).

#### 3.2.2 Identified dominant genus and species of Eukaryotes in soil (DS5) and water (DW5) samples, with several having Uranium remediation potential

Heatmap analyses of the whole-genome metagenomic dataset reveal marked differences in the relative abundance of eukaryotes in the soil (DS5) and water (DW5) samples (Figs. 16-17). In the soil sample, *Cercospora* was the predominant genus, with *Cercospora beticola* representing the most abundant species. On the other side, the water sample was dominated by the genus *Malassezia,* and the specific species was *Malassezia restricta.* This is a fungal genus, which is commonly found on animal/human skin. Furthermore, the occurrence of parasitic genera such as *Plasmodium* or *Leishmania* and the human genus *Homo sapiens* suggests environmental DNA shedding (Figs. 16-17).

**Figure 16:**
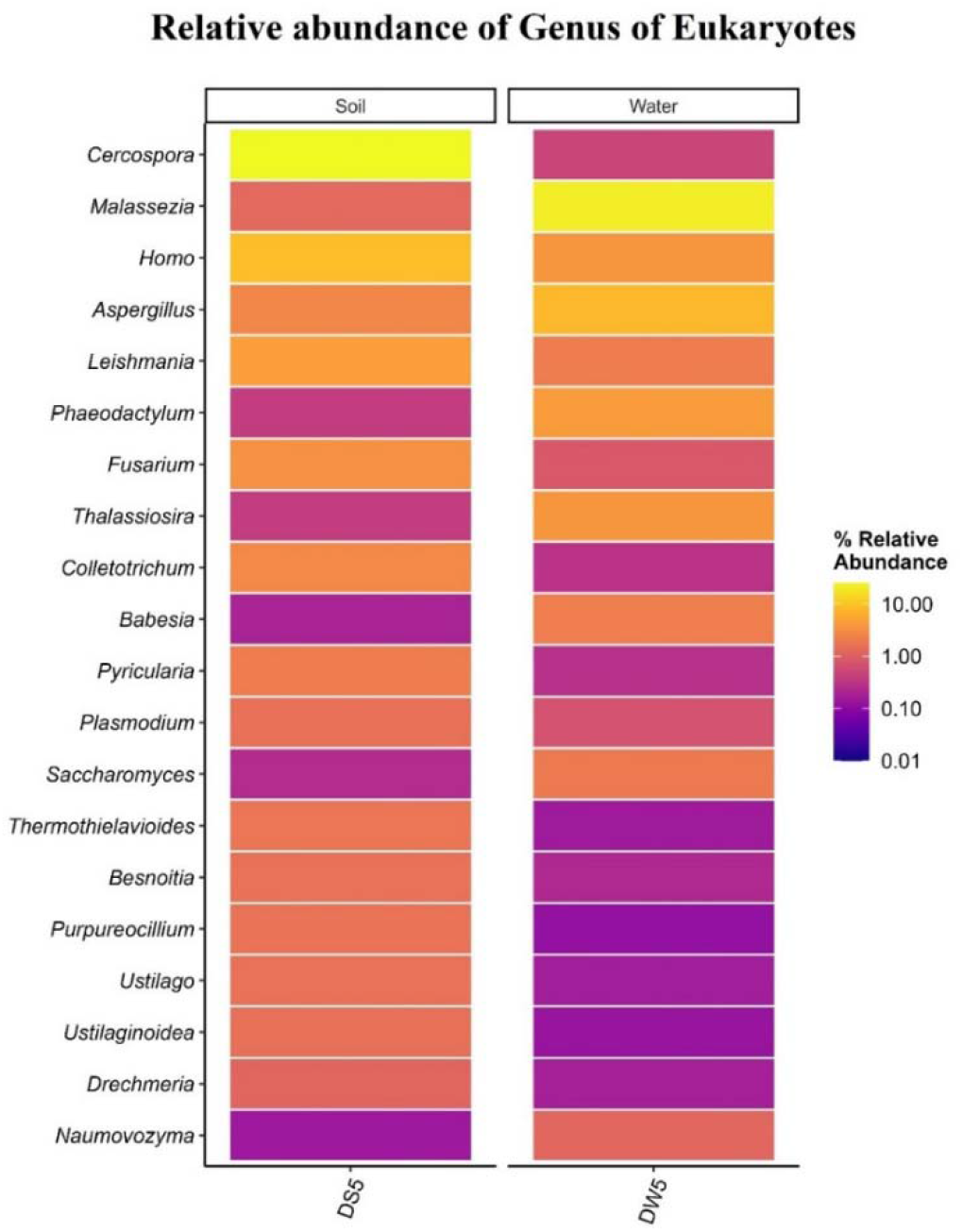
Heatmap showing the distribution of top 20 genus of Eukaryotes identified in soil (DS5) and water (DW5) samples of Dhala structure site. Each row in the heatmap represents a different genus.

**Figure 17:**
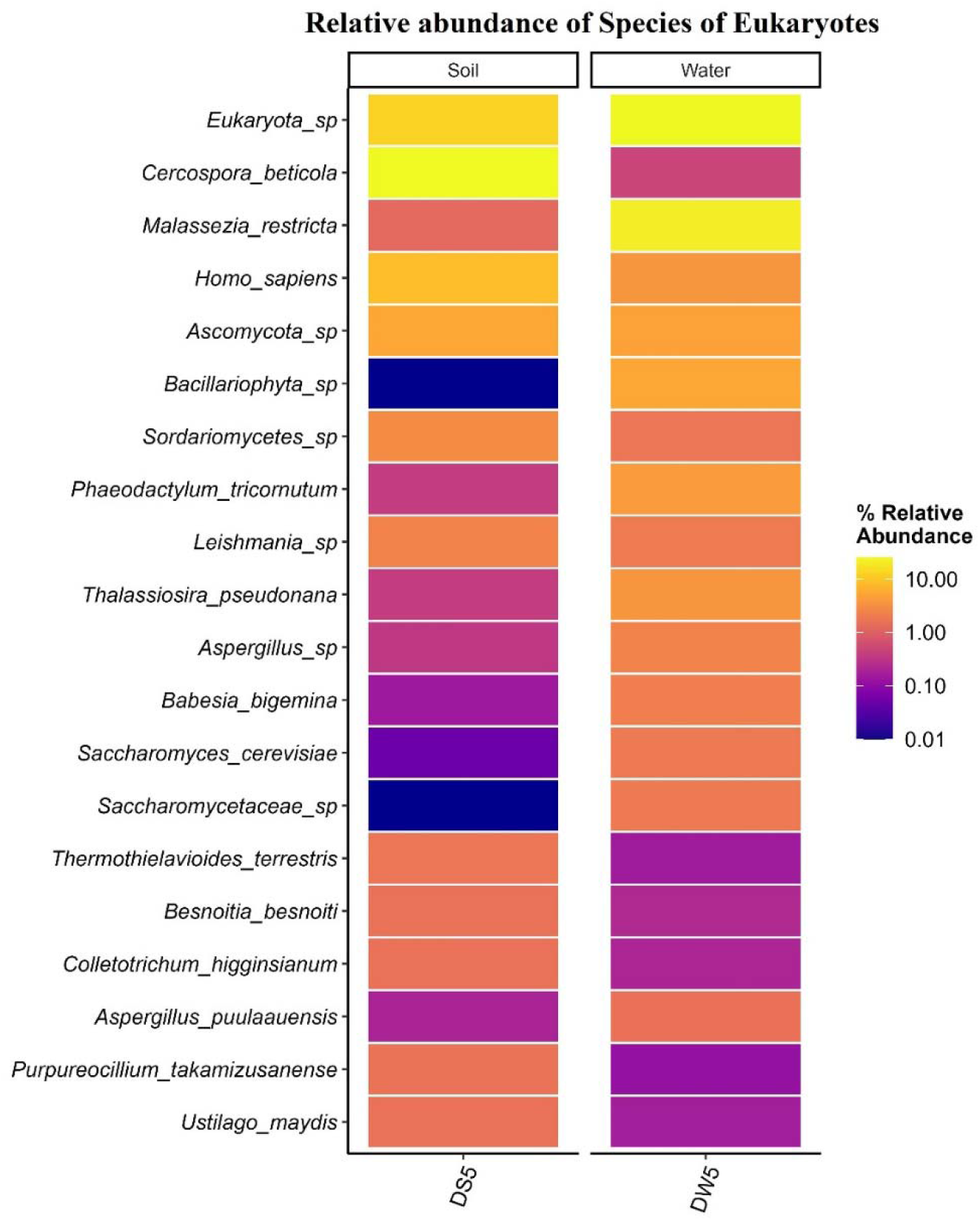
Heatmap showing the distribution of the top 20 Species of Eukaryotes identified in soil (DS5) and water (DW5) samples of the Dhala structure. Each row in the heatmap represents a different Species.

Although bacteria are widely known for bioremediation of heavy metals, the present study also identified several algal and fungal genera with well documented abilities for Uranium remediation. Genus *Aspergillus* and *Aspergillus* sp. were detected in moderate relative abundances in both the soil and water samples. *Aspergillus* spp. are well known for their roles in radionuclide and heavy metal bioremediation. *Aspergillus* spp. biomass is an excellent biosorbent for Uranium (VI), and can efficiently pull it out of aqueous habitats (Figs. 16-17).

Genus *Saccharomyces* and its specific species *Saccharomyces cerevisiae* were present in both the soil and water samples, but they exhibited higher relative abundance in the water sample. *Saccharomyces cerevisiae* is known to efficiently bind to Uranium. Apart from passive biosorption, *S. cerevisiae* cells can convert soluble Uranium into an insoluble crystalline form of Uranium termed chernikovite, thereby neutralizing its toxicity and environmental mobility (Figs. 16-17).

Another fungal genus, *Fusarium,* was detected at moderate abundance in both soil and water samples, which shows strong passive and active Uranium (VI) bioadsorption via carboxyl and hydroxyl groups present on its cell surface (Figs. 16-17).

Microalgae such as *Phaeodactylum tricornutum* and *Thalassiosira pseudonana* were identified in moderate abundance in the water sample but occurred only at relatively low abundance in the soil sample. These microalgae are well-known biosorbents for Uranium and other heavy metals via their complex cell walls, which provide many binding sites to the uranyl ions (Figs. 16-17).

### 3.3 Identification of Alpha Diversity of Viruses

The ACE index revealed that the soil sample has more viral species richness than the water sample (Fig. 18(a)). Chao1 analysis explains the total number of species in a sample. It is also helpful for the estimation of rare/unobserved species in an analyzed sample. The Chao1 analysis s revealed that the soil sample had more species richness with the presence of more rare species than the water sample (Fig. 18(b)). The coverage graph explains how well or how completely the alpha diversity of microbes has been captured in each sample by sequencing. If the coverage is >0.95, it is a very good sampling, and estimation of alpha diversity is reliable. If the coverage is <0.90, then the sampling is not good, and alpha diversity could be biased in this case. In the present study, the coverage value exceeded 0.95 for the water sample, which indicates the correct estimation of the alpha diversity of viruses. But for the soil sample, the coverage is <0.90, suggesting that the alpha diversity of viruses could be biased in this case (Fig. 18(c)). The Fisher graph basically indicates viral diversity in the given sample. In the present Fisher graph, a very high diversity of viral populations was observed for the soil than the water sample, suggesting high species richness, more rare taxa, and the possibility of a more complex viral community (Fig. 18(d)). The Inverse Simpson (InvSimpson) graph mainly helps in finding evenness and dominance of taxa in a viral community, or in other words, it tells how balanced your viral community. If there is a high InvSimpson value that means there is a presence of various viruses, all are present in the same number, and no one is dominating. Therefore, it can be termed as a balanced and healthy community. The InvSimpson index was markedly higher in the soil sample, indicating a more even distribution of viral taxa and a comparatively balanced community structure. Conversely, the lower Inverse Simpson value observed in the water sample suggests a limited number of highly abundant viral taxa (Fig. 18e).The observed graph reveals about how many different types of viruses are present in any sample. Water sample show a high bar value that means it contains various types of viruses. Also, there is a high richness and complex viral community. Whereas, low value in the soil sample suggested the presence of fewer viruses, low richness, and a disturbed environment (Fig. 18(f)). Pielou’s evenness index plot represents how balanced or even the distribution of species is. In the present study, we constructed this plot to study the distribution pattern of viruses. In soil sample, the value is >0.5, there is a moderate imbalance among viral species. Whereas in the water sample, the value is <0.5, which means there is dominance of a few taxa (Fig. 19(a)). The Shannon graph tells about the species diversity and richness. The water sample shows moderate diversity of viruses. Whereas the soil sample shows a high diversity of viruses (Fig. 19(b)). In a Simpson graph, higher values indicate high diversity, and lower values indicate low diversity. In the analyzed water sample, the value was low, so there is low diversity of viruses. Whereas, in the soil sample, there is a very high value, indicating a high diversity of viruses (Fig. 19(c)).

**Figure 18:**
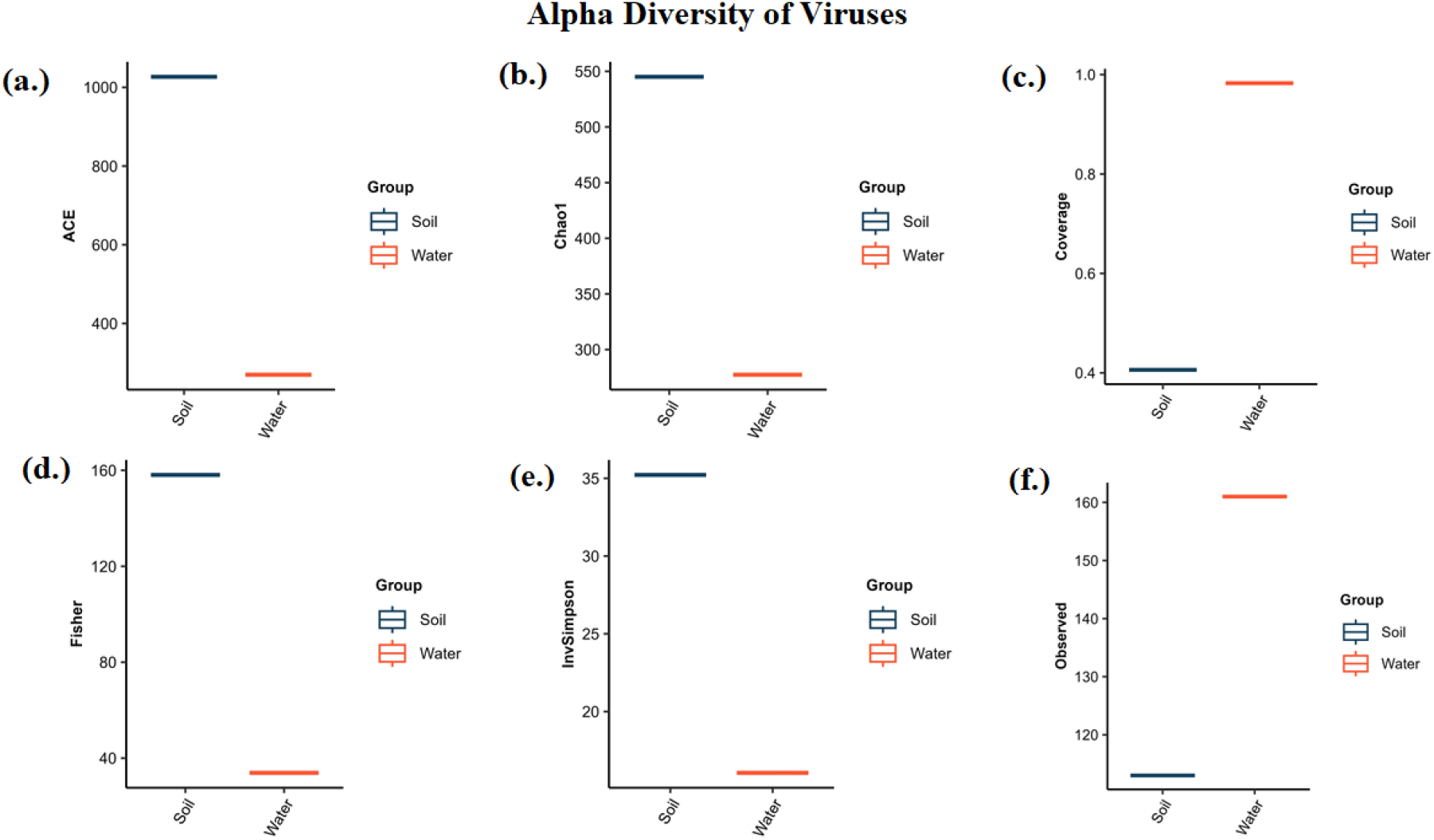
Alpha diversity analysis of the Viral population present in the soil and water sample collected from the Dhala area; **(a.)** ACE, **(b.)** Chao1 **(c.)** Coverage **(d.)** Fisher **(e.)** InvSimpson **(f.)** Observed.

**Figure 19:**
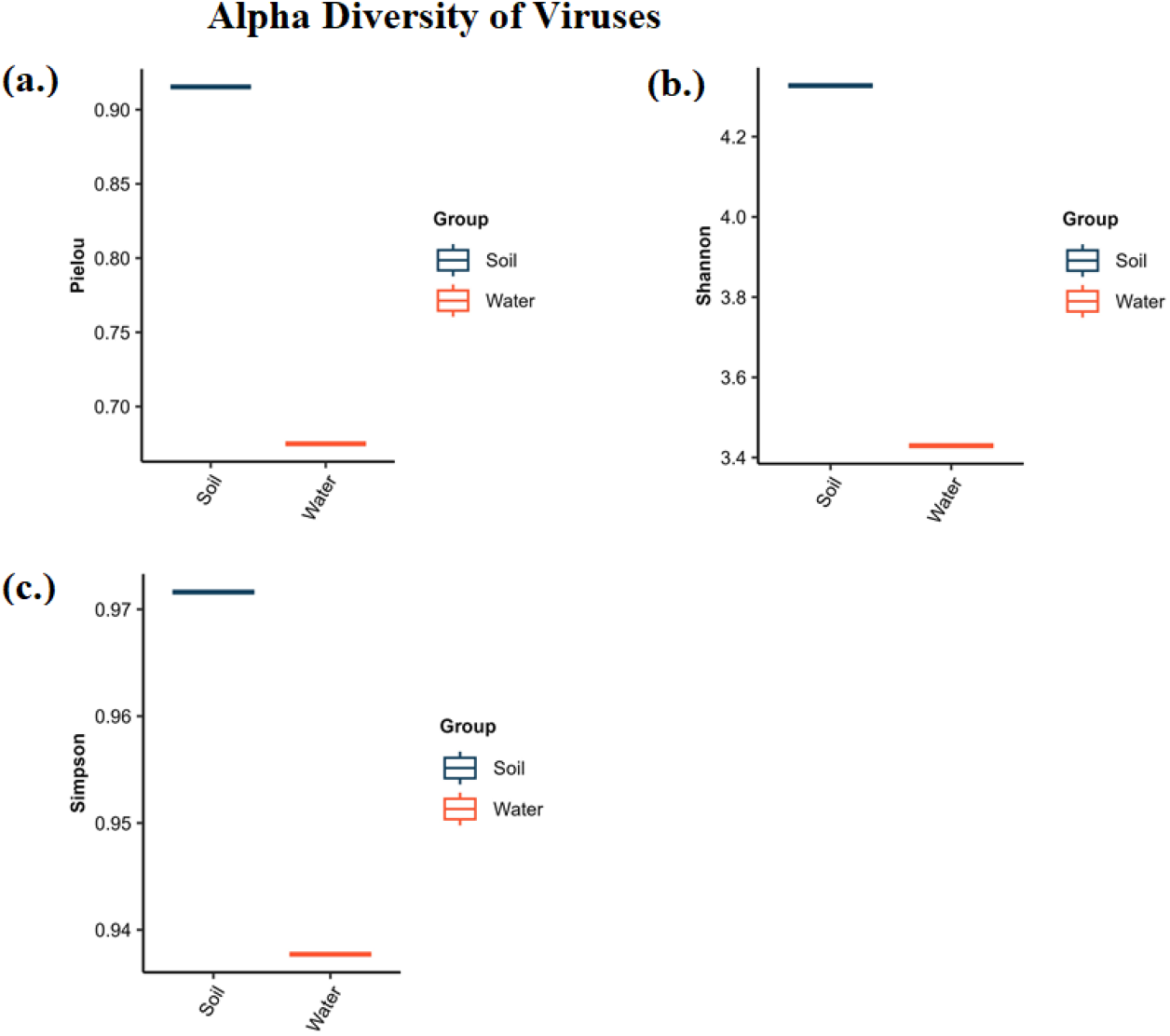
Alpha diversity analysis of the Viral population present in the soil and water sample collected from Dhala; **(a.)** Pielou, **(b.)** Shannon, **(c.)** Simpson.

#### 3.3.1 Identification of Relative Abundance of Phylum, Class, Order, Family, Genus, and Species of Viruses in the Analyzed Soil and Water Samples

During phylum identification, in the soil sample, phylum *Uroviricota* was abundant. Whereas, *Prploviricota, Nuleocytoviricota, Saleviricota, Cossaviricota, Pisuviricota,* and *Kitrinoviricota* were less abundant. In the water sample, *Uroviricota* was abundant. While *Phixviricota, Pisuviricota, Saleviricota,* and *Hofneiviricota* were less abundant (Fig. 20 (a,b)). During the identification of the class in the soil sample, *Caudoviricetes* were abundant. Whereas, *Herviricetes, Naldaviricetes, Pokkesviricetes, Megaviricetes, Papovaviricetes, Pisoniviricetes,* and *Alsuviricetes* were less abundant. In the water sample, *Caudoviricetes* were abundant. Whereas, *Malgrandaviricetes, Megaviricetes,* and *Huolimaviricetes* were less abundant (Fig. 20 (a,b)).

**Figure 20:**
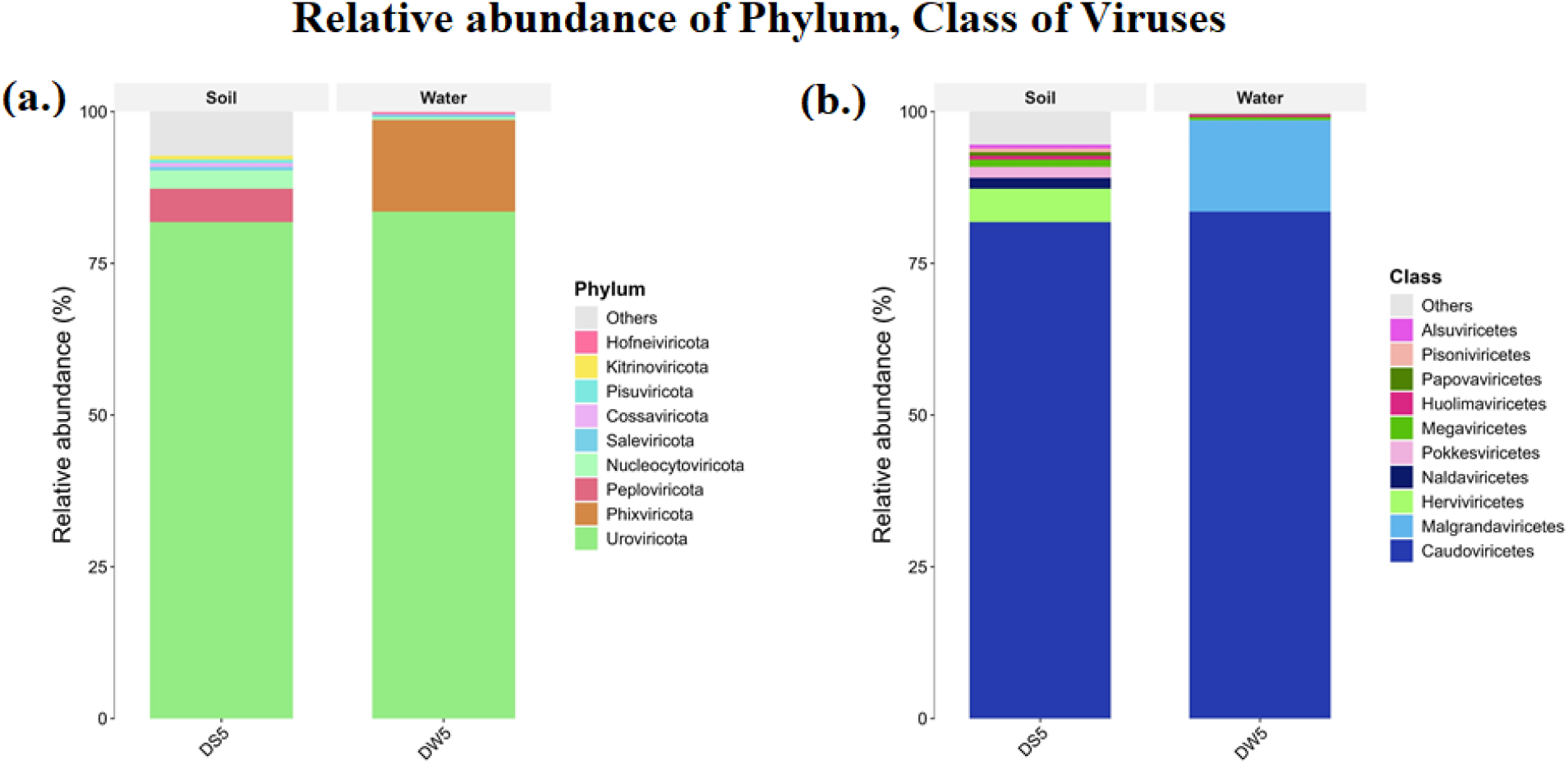
Stack bar plot showing the relative abundance of Phylum, Class of Viruses

During order identification, in the soil sample, *Herpesvirales* were abundant. While *Lefavirales, Chitovirales, Imitervirales, Zurhausenvirales, Nidovirales, Thumleimavirales,* and *Martellivirales* were less abundant. In the water sample, *Petitvirales* were abundant. Whereas *Thumleimavirales,* and *Haloruvirales* were less abundant. In the family identification of soil sample, *Herpesviridae,* and *Pseudoviridae*were abundant. Whereas *Herelleviridae, Hafunaviridae, Baculoviridae, Casjensviridae,* and *Poxviridae*. In the water sample, families such as *Straboviridae, Microviridae,* and *Herelleviridae* were found to be abundant. Whereas *Mesyanzhinovviridae* and *Hafunaviridae* were less abundant families (Fig. 21 (a,b)).

**Figure 21:**
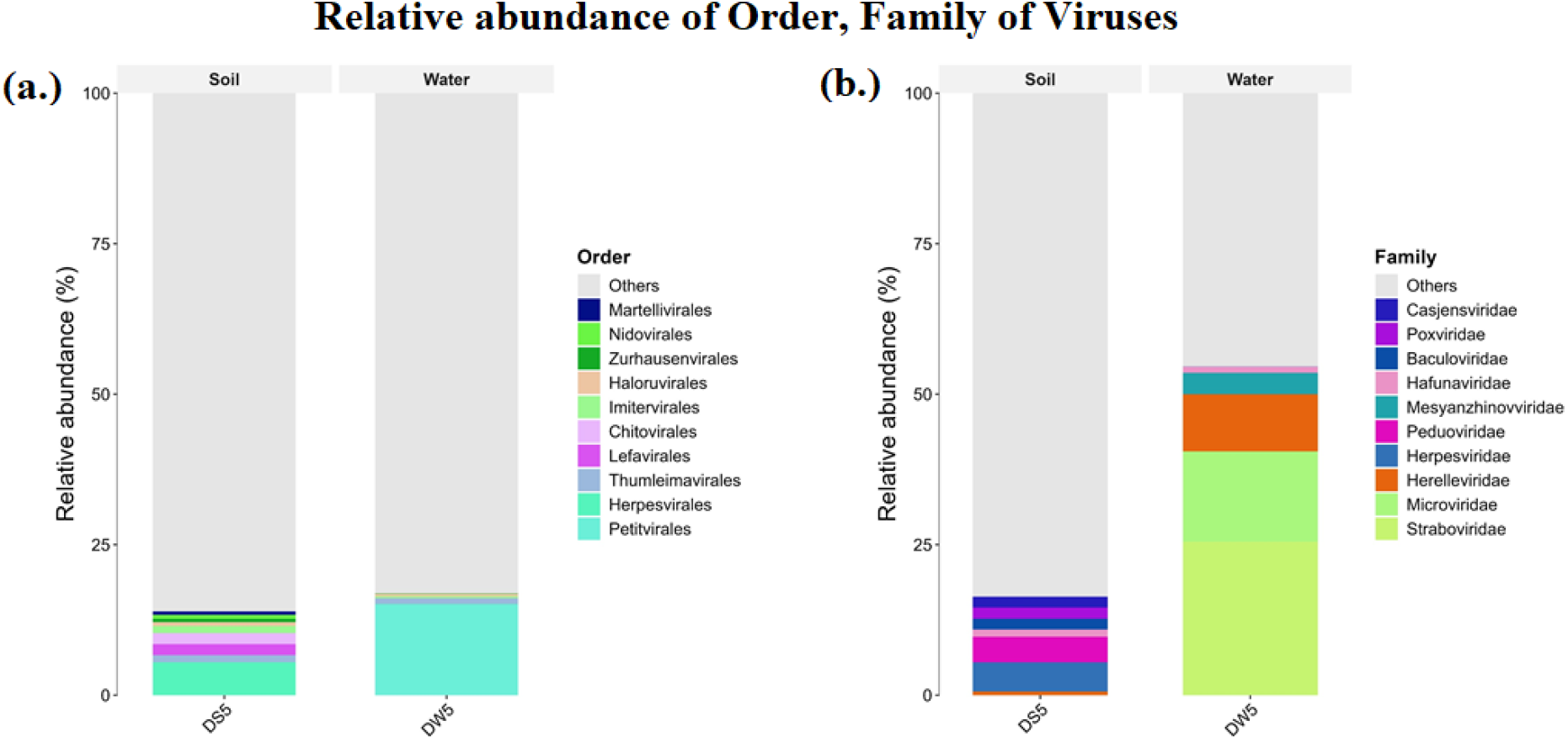
Stack bar plot showing the relative abundance of Order, Family of Viruses

During Genus identification, in the soil sample, *Pandovirus, Bielevirus, Seussvirus, Colossusvirus,* and *Simplexvirus* were abundant, whereas *Justusliebigvirus* was less abundant. In the water sample, *Tulanevirus, Moineauvirus,* and *Siminovitchvirus* were abundant. Whereas *Justusliebigvirus* and *Unaquatrovirus* were less abundant (Fig. 22 (a,b)).

**Figure 22:**
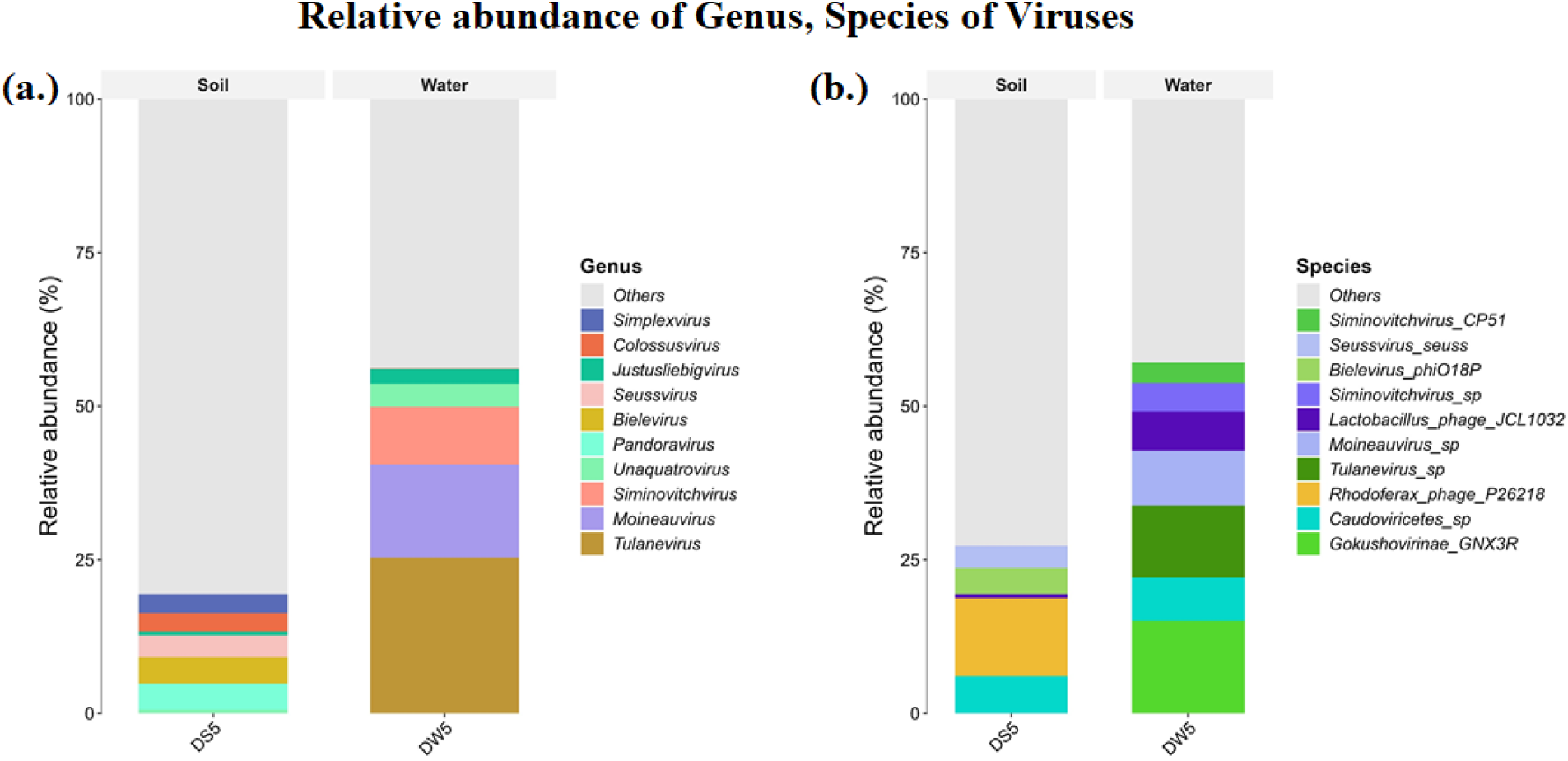
Stack bar plot showing the relative abundance of Genus, Species of Viruses

During species identification in the soil sample, *Rhodoferax_phage_P26218,* and *Caudoviricetes_sp* were abundant, whereas *Bielevirus_phiO18P, Seussvirus_seuss,* and *Lactobacillus_phage_JCL1032* were less abundant. In the water sample, *Gokushovirinae_GNX3R, Tulanevirus_sp, Moineauvirus_sp,* and *Caudoviricetes_sp* were abundant, whereas *Lactobacillus_phage_JCL1032, Siminovitchvirus_sp,* and *Siminovitchvirus_CP51* were less abundant (Fig. 22 (a,b)).

#### 3.3.2 Identified dominant genus and species of Viruses in soil (DS5) and water (DW5) samples, with their role in Uranium remediation

In the soil sample, genera such as *Pandoravirus, Bielevirus, Simplexvirus, Seunavirus,* and *Colossusvirus* were found in abundance. At the specific species level, *Rhodoferax phage P26218* and *Caudoviricetes* sp. were abundant in the soil sample (Fig. 23,24).

**Figure 23:**
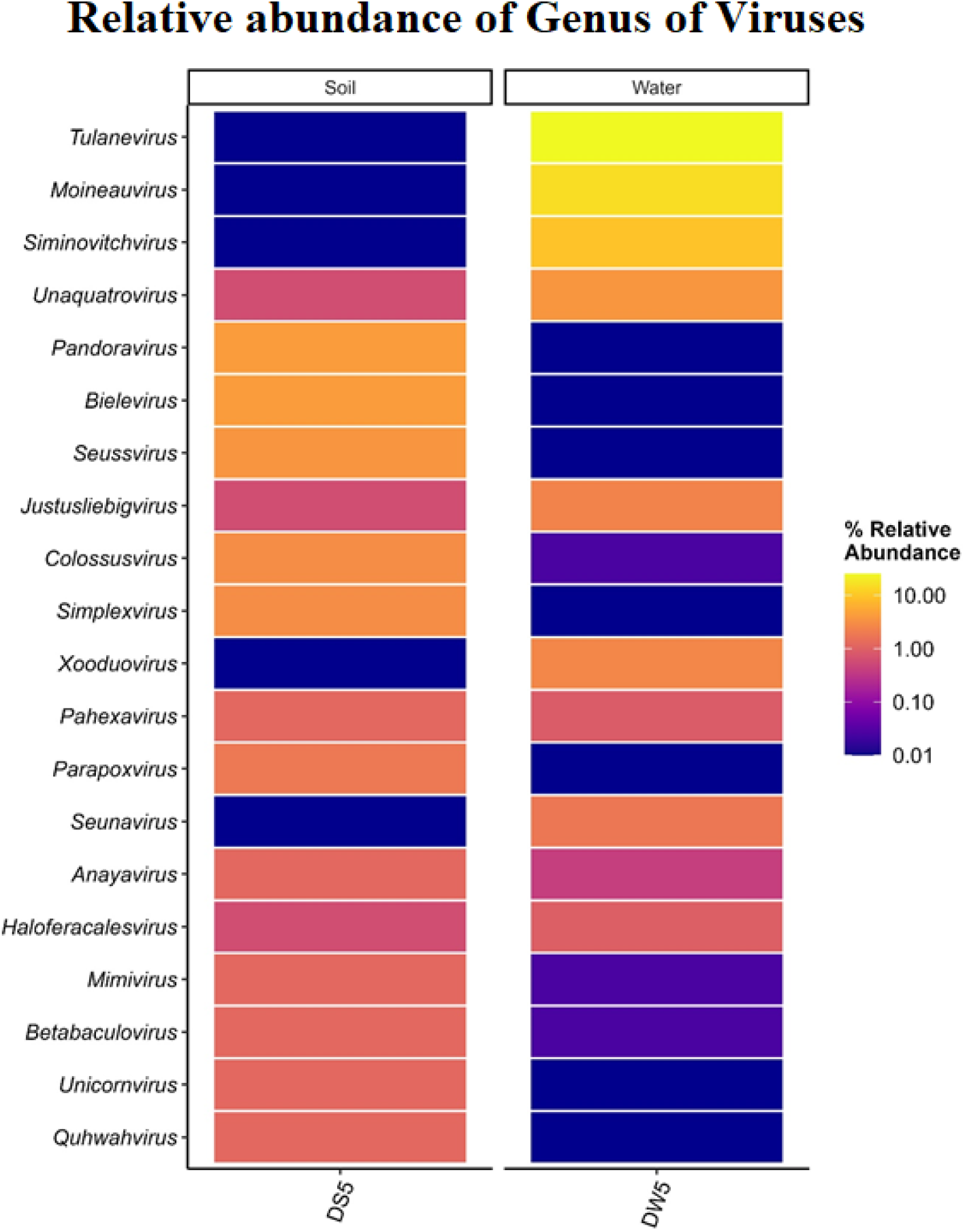
Heatmap showing the distribution of the top 20 Genus of Viruses identified in soil (DS5) and water (DW5) samples from the Dhala structure. Each row in the heatmap represents a different Genus.

**Figure 24:**
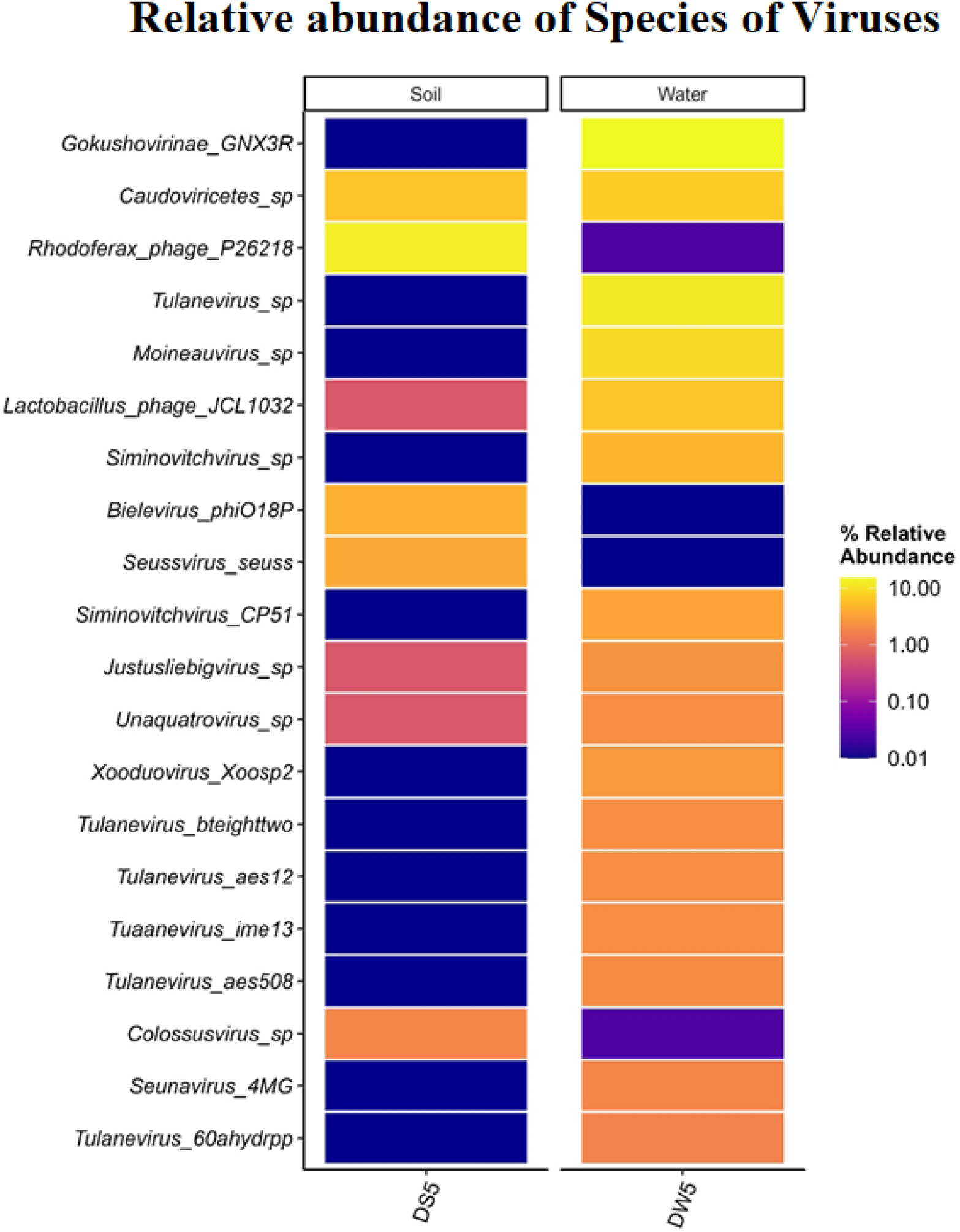
Heatmap showing the distribution of the top 20 Species of Viruses identified in soil (DS5) and water (DW5) samples from the Dhala structure. Each row in the heatmap represents a different Species.

In the water sample, genera *Tulanevirus, Siminovitchvirus,* and *Moineauvirus* were found to be highly abundant. At the specific species level, *Tulanevirus* sp., *Tulanevirus bteighttwo, Siminovitchvirus CP51,* and *Moineauvirus* sp. were found to be abundant in the water sample. Apart from these species, the *Gokushovirinae GNX3R* species was also found in abundance in the water sample (Fig. 23,24).

Viruses can not remediate the Uranium. The abundance of *Rhodoferax phage P26218* in the soil sample suggests the presence of *Rhodoferax* bacteria. This bacterium is well known for its capability to reduce heavy metals and iron and finally perform the process of bioremediation. *Rhodoferax phage P26218* regulates the population of *Rhodoferax* bacteria by infecting and lysing it. This process influences the overall success and rate of Uranium and heavy metal reduction in that particular environment (Fig. 23,24).

### 3.4 Common and unique taxa between soil and water samples

Venn diagrams were constructed to estimate common and unique taxa in soil and water samples (Fig. 25). Among bacteria and archaea, 6249 taxa were common to both habitats, 1474 were exclusively detected in the soil sample and 895 unique taxa were found in the water sample, respectively (Fig. 25(a)). Among eukaryotes, a total of 101 taxa were common in both the soil and water samples, while 25 and 29 taxa were unique to the soil and water samples, respectively (Fig. 25(b)). Among viruses, only 14 taxa were common to both the soil and water samples, whereas 99 and 147 taxa were uniquely associated with the soil and water samples, respectively (Fig. 25(c)).

**Figure 25:**
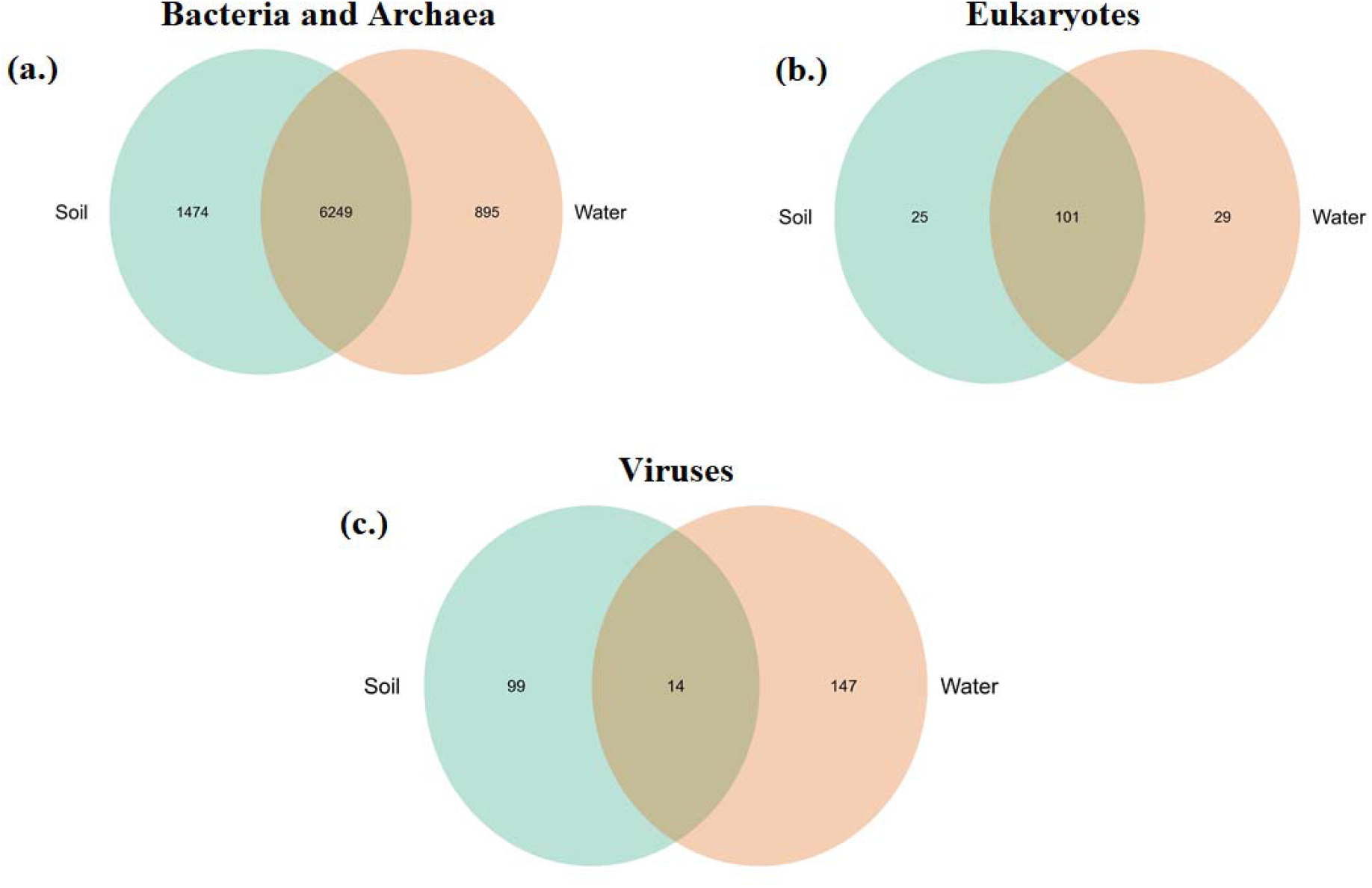
Venn diagrams depicting common and unique taxa between soil and water samples from the Dhala area; **(a.)** Bacteria and Archaea **(b.)** Eukaryotes, and **(c.)** Viruses.

### 3.5 Functional Attributes of a Uranium-Rich Site (Impact Melt Breccia) in the Dhala Impact Structure

Whole metagenomic sequencing of the soil and water samples from the Dhala area identified 58 genes responsible for extracellular electron transfer, metal tolerance, and environmental stress response. This suggests that microbial communities in both soil and water possess a genetic repository, possessing a crucial role in uranium bioremediation.

The copA gene, which encodes a P-type Cu^+^ transporter involved in heavy metal efflux, was found in both samples, with 399 reads in soil and 432 reads in water. Genes czcA encodes a heavy metal efflux system, and the chrA chromate transporter were enriched in the water sample. Gene czcA gene was represented by 97 reads in soil, and 273 reads in water, whereas chrA recorded 143 and 308 reads, respectively. The high abundance of these efflux pumps suggests that the microbes present in the studied site are defending themselves against uranium and many other heavy metals present in the environment. The presence of czcA complex functions to efflux zinc, cobalt, and cadmium. Furthermore, the presence of chromate and copper transporters indicates that remediation of uranium is performed by microbes present in the studied site. The above findings suggest that the microbiota present in this habitat have the ability to survive heavy metal toxicity and can help in the reduction of uranium (VI) (Tables 1-2 & Fig. 26).

**Figure 26:**
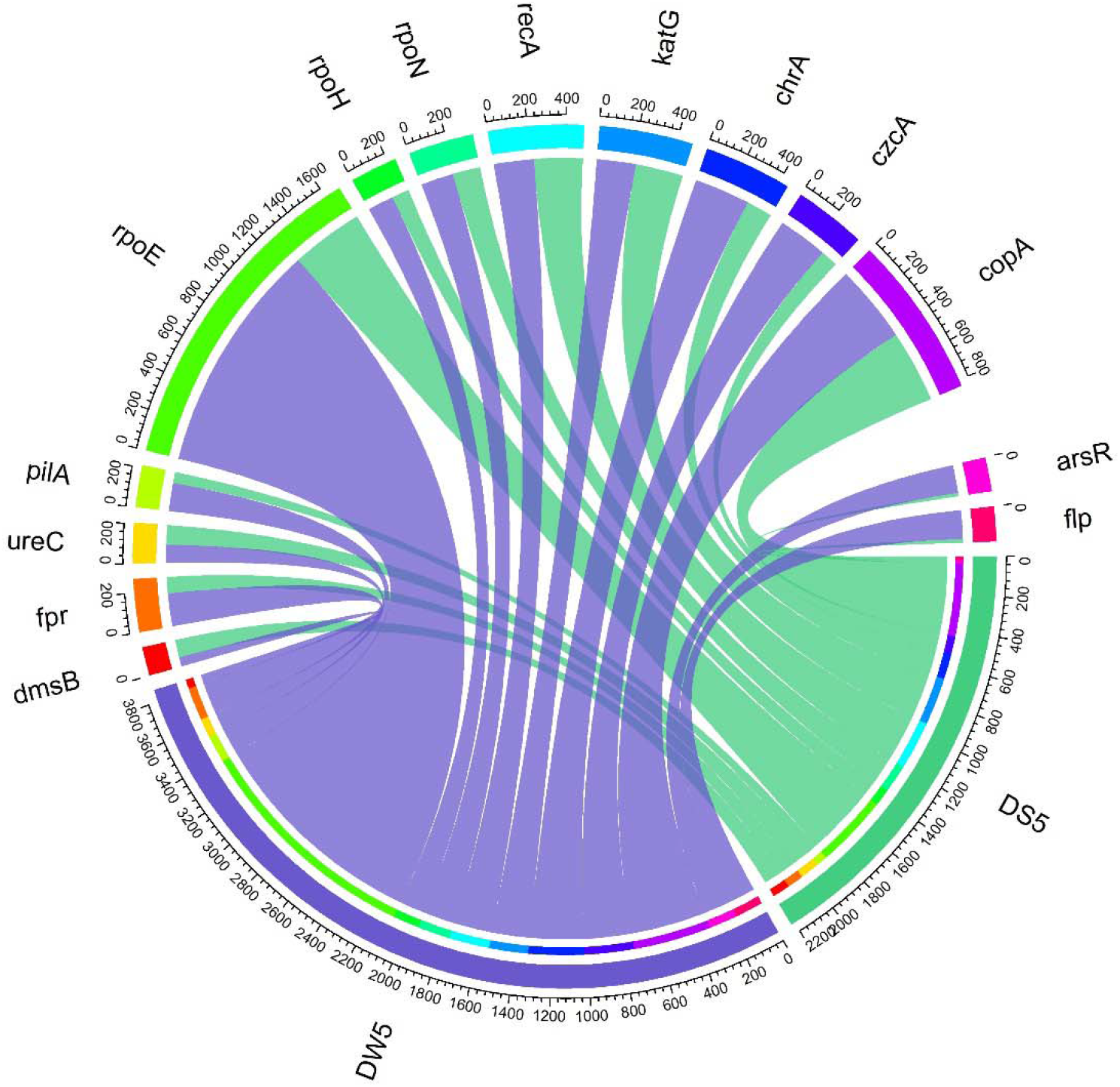
Distribution of the top 14 uranium bio-remediation genes in soil (DS5) and water (DW5) samples visualized by CIRCOS software.

**Table 1:**
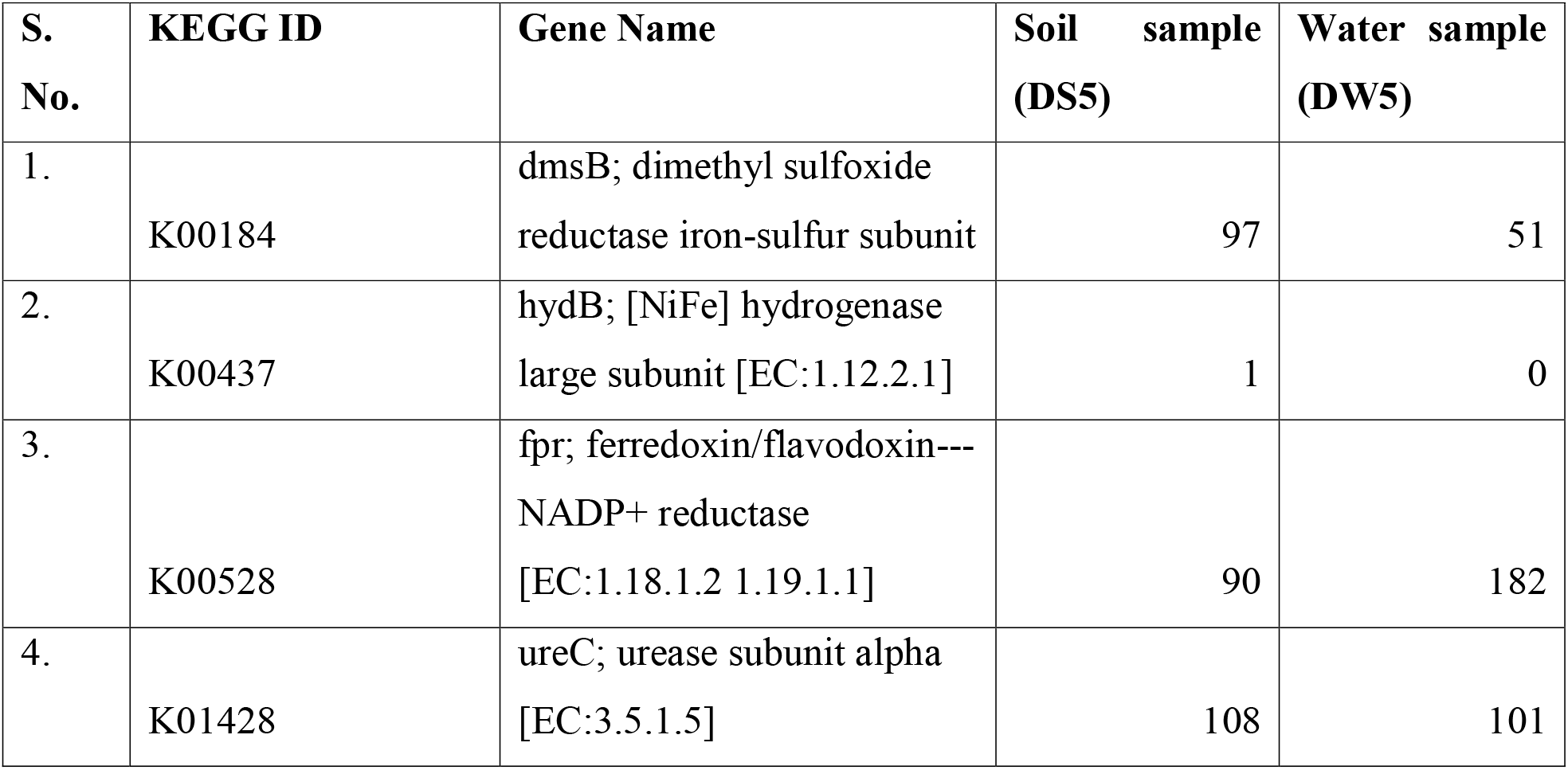
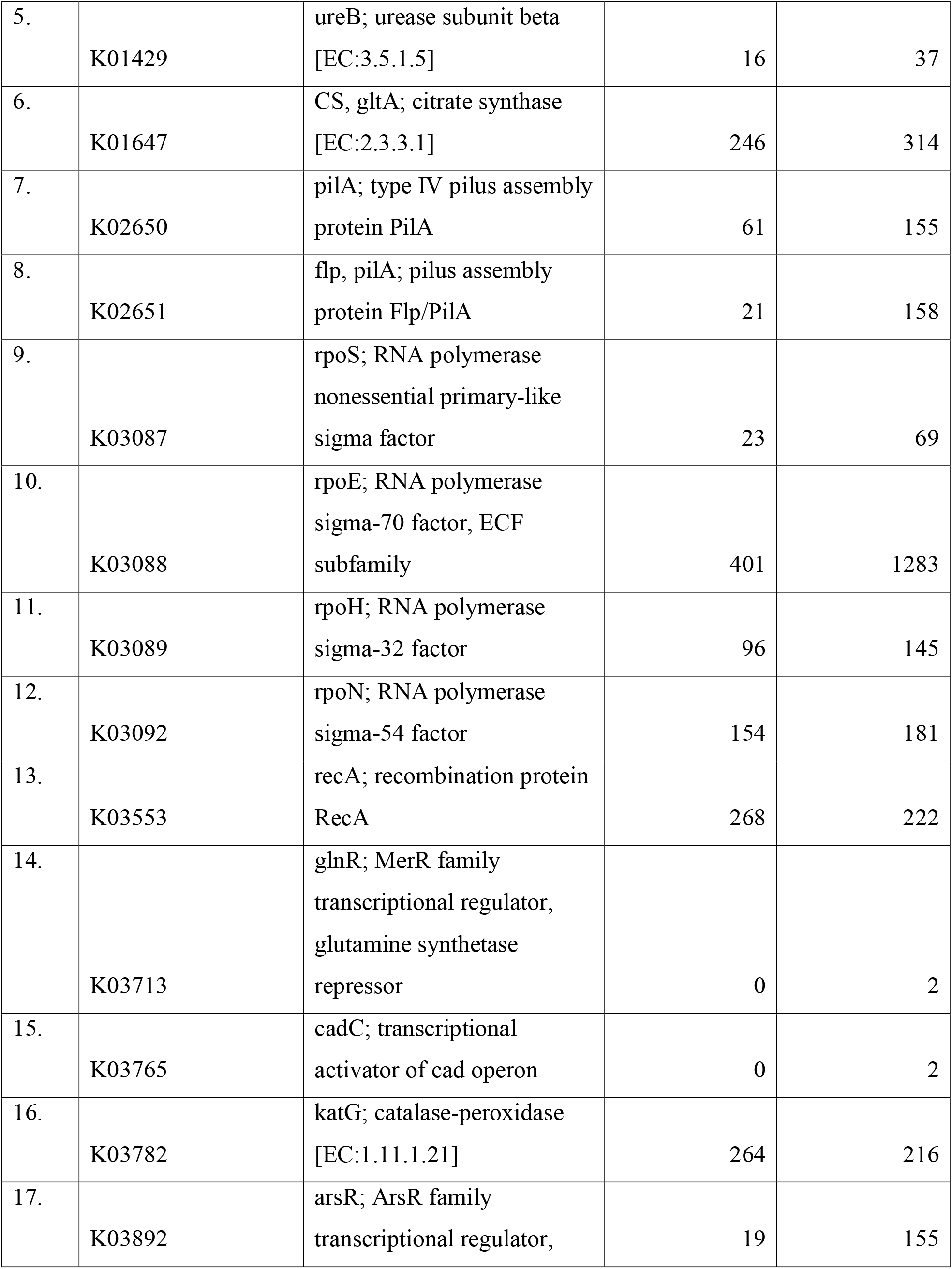
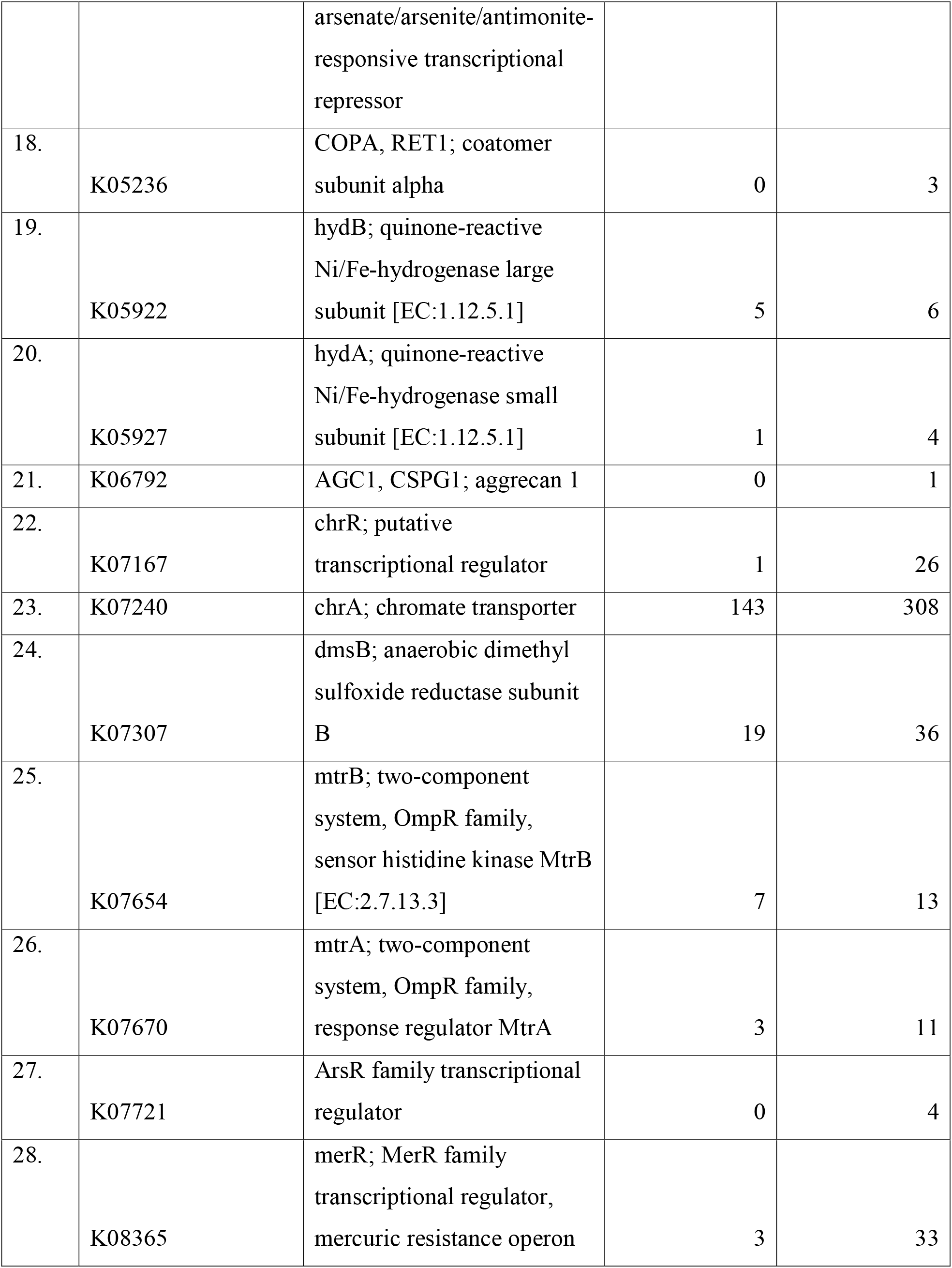
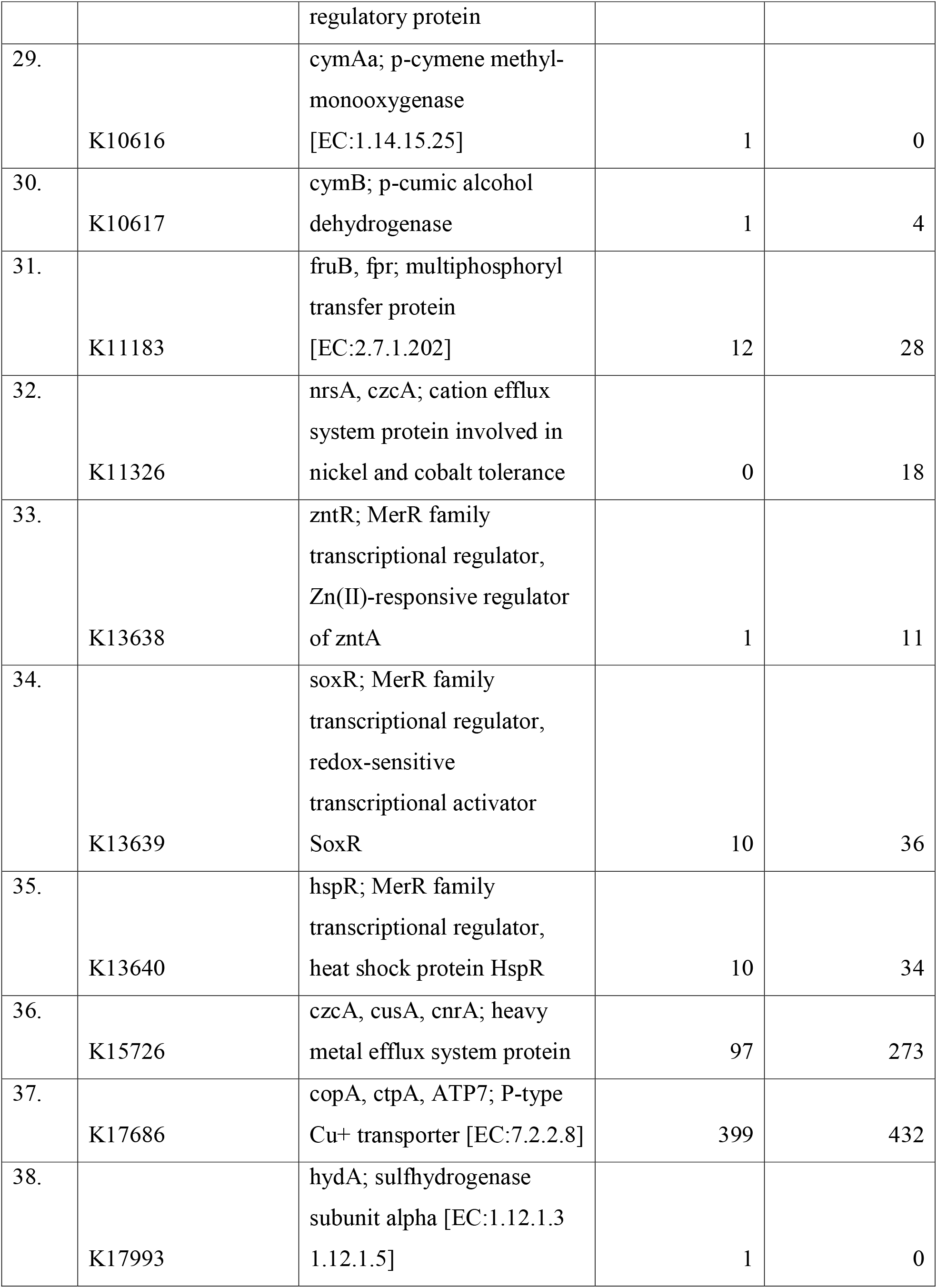
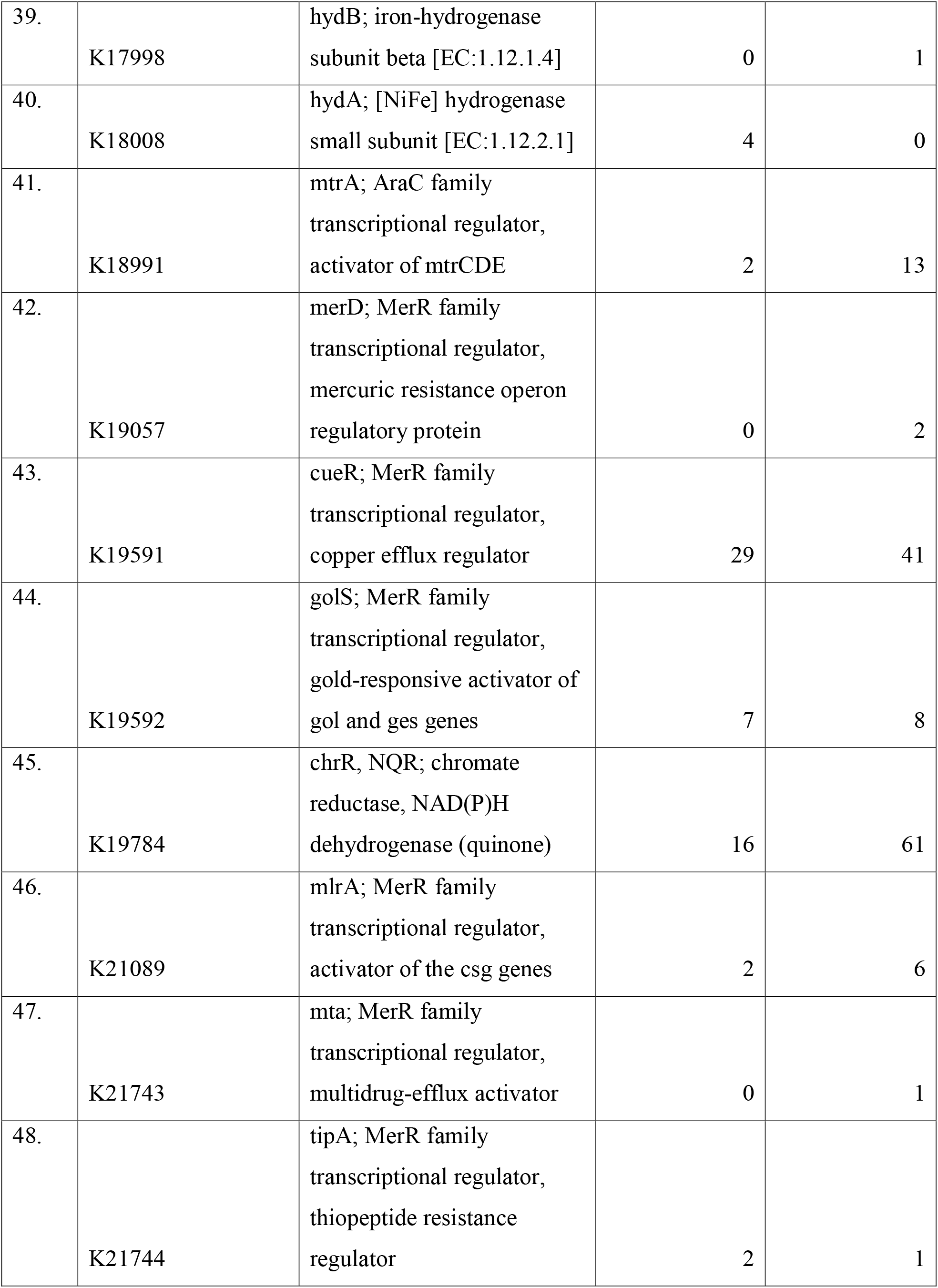
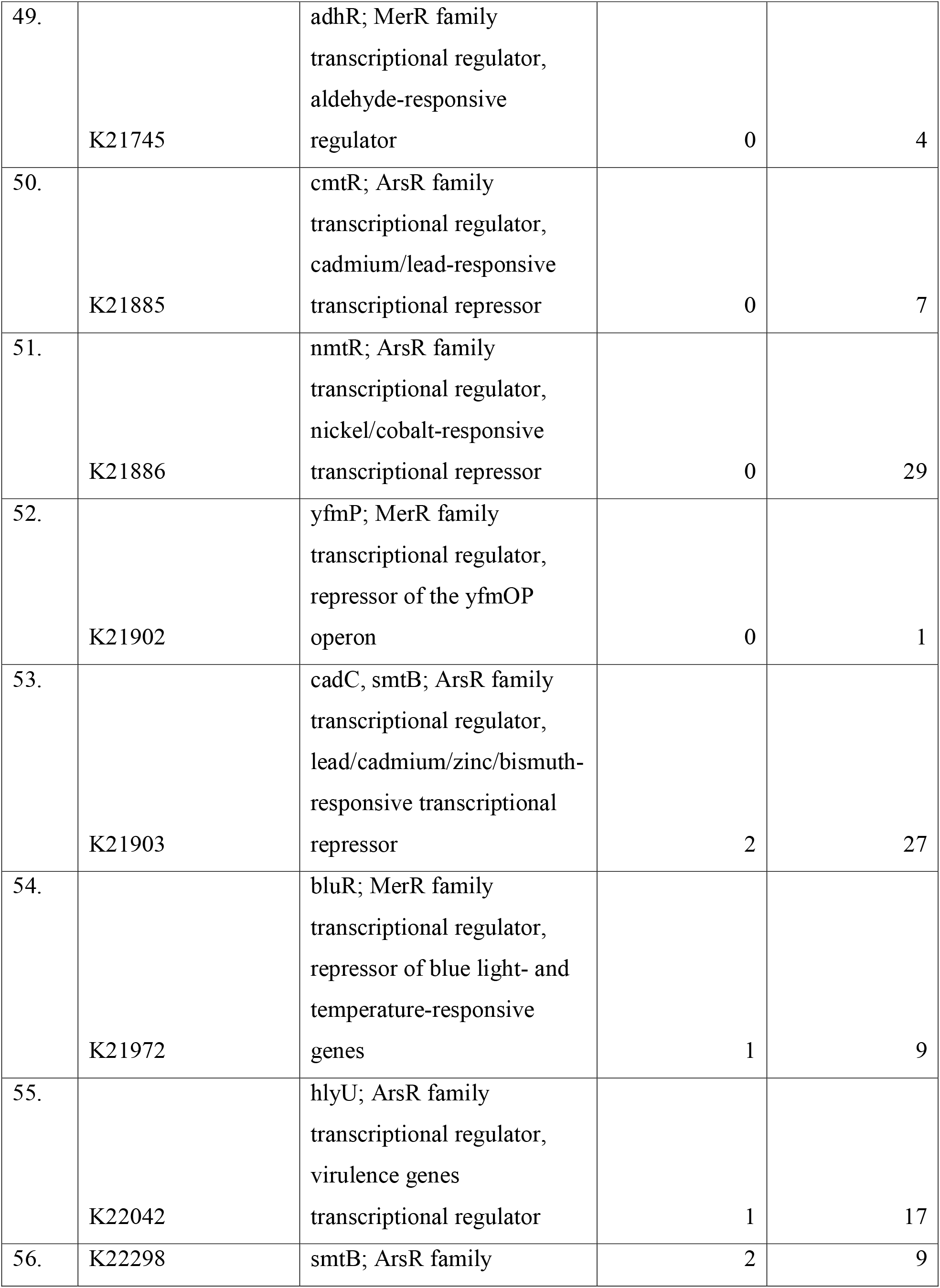
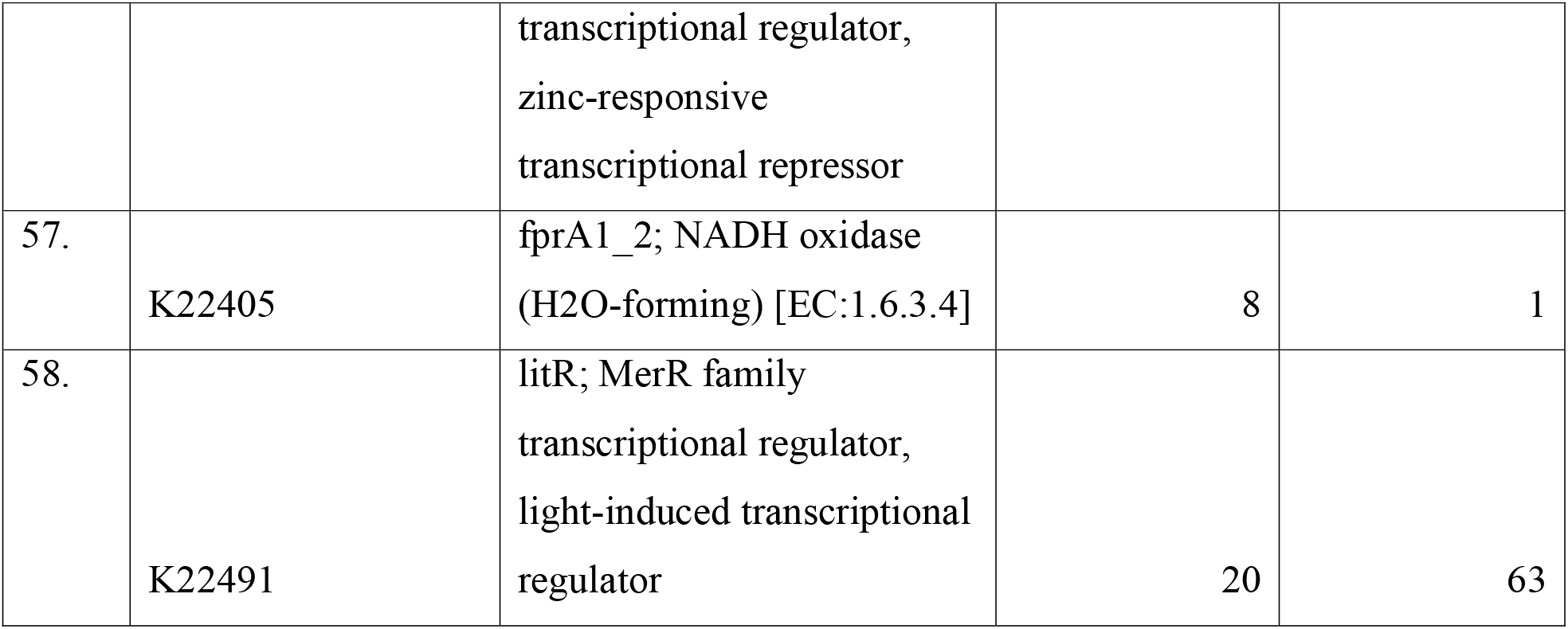
Uranium bio-remediation genes detected in soil (DS5) and water (DW5) samples from the western part of the Dhala structure, India.

**Table 2:** Top 14 uranium bio-remediation genes detected in soil and water samples collected from the western part of the Dhala structure, India.

| S. No. | Gene name | Soil sample (DS5) | Water sample (DW5) |
| --- | --- | --- | --- |
| 1. | dmsB | 97 | 51 |
| 2. | fpr | 90 | 182 |
| 3. | ureC | 108 | 101 |
| 4. | pilA | 61 | 155 |
| 5. | rpoE | 401 | 1283 |
| 6. | rpoH | 96 | 145 |
| 7. | rpoN | 154 | 181 |
| 8. | recA | 268 | 222 |
| 9. | katG | 264 | 216 |
| 10. | chrA | 143 | 308 |
| 11. | czcA | 97 | 273 |
| 12. | copA | 399 | 432 |
| 13. | arsR | 19 | 155 |
| 14. | flp | 21 | 158 |

In both samples, gene such as dmsB (dimethyl sulfoxide reductase iron-sulfur subunit) for the enzymatic reduction of heavy metals was detected , with 97 reads in the soil sample and 51 reads in the water sample, respectively. In addition, the pilA type IV assembly proteins (K02650 and K02651) were abundant in the water sample, yielding 155 and 158 reads, respectively compared with 61 and 21 reads in the soil sample (Tables 1-2 & Fig. 26).

Uranium bioremediation takes place through the enzymatic reduction of toxic soluble U(VI) to inert insoluble U(IV) form. The abundance of pilA, type IV pilus assembly proteins in the water sample indicates potential for extracellular electron transfer. Type IV pili acts as ‘nanowires’, which facilitates the direct transport of electrons U(VI) (extracellular receptor). The identification of trace hydrogenases (hydA and hydB), and reductases dmsB further revealed a functional metabolic network of biological shuttles and electron donors having the potential to perform the reduction of heavy metals in the studied habitats. Metagenomic insight further revealed an intense regulatory response to environmental stresses. The most abundant gene in the whole dataset was rpoE (RNA polymerase sigma factor) represented by 1283 reads in the water sample and 401 reads in the soil sample. Similarly, katG, an oxidative stress marker (catalase-peroxidase), was also in high abundance in both water (216 reads) and soil (264 reads). Metal-responsive transcriptional regulators belonging to MerR and ArsR were detected in the studied samples. The transcriptional repressor arsR exhibited a higher abundance in the water sample (155 reads) than in the soil sample (19) (Tables1-2 & Fig. 26).

In a nutshell, metagenomics insights into the soil and water samples from the Dhala area revealed the potential of microbial populations for uranium bioremediation. The identification of genes responsible for extracellular electron transfer, such as pilA (type IV pilus assembly proteins) and dmsB (reductases), indicates a functional biological pathway having the potential to reduce toxic soluble U (VI) into a stable, insoluble U(IV). Further, czcA, copA, chrA (metal efflux complexes), and rpoE (extreme stress response sigma factors) together revealed the extraordinary physiological resilience. The extremophilic communities identified in the present whole genome metagenomics study are naturally evolved and genetically well equipped to tolerate high multi-metal toxicity, and provide a self-sustaining and highly effective mechanism for environmental cleanup (Tables 1-2 & Fig. 26).

## 4. Discussion

### 4.1 Cross-domain microbial diversity in the soil and water samples from the Dhala area

The present whole-genome metagenomic investigation demonstrates that the Dhala structure hosts a remarkably diverse cross-domain microbiome that has undergone long-term adaptation to a naturally uranium-enriched impact-affected environment. The dominance of *Pseudomonadota*, *Actinomycetota*, and *Cyanobacteriota*, together with diverse archaeal, fungal, and viral communities, indicates that the investigated samples from the Dhala area provide a wide range of ecological niches supporting microorganisms with complementary metabolic strategies [4,5,34,35]. Unlike microbial communities inhabiting homogeneous terrestrial ecosystems, the Dhala microbiome reflects the combined influence of impact-induced fracturing, hydrothermal alteration, prolonged weathering, groundwater circulation, and a natural radionuclide-enriched geological setting [4,12,2,3,14]. These geological processes have collectively generated a metabolically heterogeneous ecosystem capable of sustaining microorganisms possessing broad physiological plasticity and extensive functional diversity [3,34,14].

The contrasting diversity patterns observed between the soil and water samples from the Dhala area further emphasize the importance of habitat heterogeneity in microbial community assembly. Such partitioning of microbial diversity likely enhances ecosystem resilience by increasing functional redundancy while allowing specialized microorganisms to occupy distinct physicochemical niches [36,34,37].

### 4.2 Comparison with uranium mine, impact crater and deep biosphere microbiomes

The taxonomic and functional characteristics of the Dhala microbiome exhibit several similarities to microbial communities previously reported from uranium mines, mill tailings, and radionuclide-contaminated environments [11,12,14]. Dominant bacterial genera including *Pseudomonas, Burkholderia, Sphingomonas*, and other proteobacterial lineages have frequently been associated with uranium-rich environments because of their exceptional metabolic versatility, efficient metal transport systems, and tolerance to oxidative stress [12,38]. The enrichment of genes encoding metal efflux pumps, ATP-dependent transporters, oxidative stress enzymes, hydrogenases, and DNA repair proteins represents a common genomic signature of microorganisms inhabiting metal-contaminated ecosystems [13,14]. However, despite these similarities, the Dhala microbiome differs fundamentally from anthropogenically contaminated uranium sites as the mine environments represent relatively recent ecological disturbances, but the Dhala structure has experienced nearly two billion years of natural geological evolution. As a result, the indigenous microbiome likely reflects long-term evolutionary adaptation rather than short-term ecological selection. This distinction provides an exceptional opportunity to investigate naturally evolved mechanisms of radionuclide resistance and microbial ecosystem stability under prolonged geochemical stress [14,39].

Microbial communities inhabiting terrestrial impact structures are increasingly recognized as valuable analogues of deep subsurface ecosystems, as impact-generated fracturing enhances groundwater circulation, water-rock interaction, and promotes long-term habitat connectivity [4,5, 6,7,2,3,40]. Similar to observations from Chicxulub structure, Chesapeake Bay structure, and other impact-generated subsurface environments, the Dhala microbiome is also dominated by metabolically versatile microorganisms capable of exploiting mineral-derived energy sources under oligotrophic conditions. The prevalence of *Pseudomonas, Sphingomonas*, and other metabolically flexible Proteobacteria in the Dhala samples further suggests adaptation to nutrient-limited conditions through broad catabolic capabilities, efficient metal homeostasis, and diverse respiratory pathways that enable survival under fluctuating redox conditions [13,12,14]. The presence of chemolithotrophic bacteria, metabolizing archaea, and mineral-associated fungi indicates that multiple trophic strategies coexist within the impact-generated lithologies.

The microbial community structure observed at Dhala also exhibits remarkable similarities to those reported from deep crystalline bedrock, Precambrian shield environments, and continental deep biosphere ecosystems. These habitats are typically characterized by extreme nutrient limitation, low rates of energy turnover, restricted organic carbon availability, and long-term geological stability, conditions that strongly select for microorganisms possessing efficient stress-response systems, slow but flexible metabolic networks, and extensive physiological plasticity [41,42,43,44]. In agreement with these studies, functional metagenomic analysis of the Dhala microbiome revealed extensive enrichment of genes associated with membrane transport, ATP-dependent metal efflux, DNA repair, oxidative stress protection, sulfur metabolism, hydrogen utilization, and extracellular electron transfer. These metabolic functions represent hallmark genomic adaptations that enable microbial survival under persistent metal toxicity, oxidative stress, and energy-limited conditions commonly encountered in the continental deep biosphere [13,12,44,14].

### 4.3 Environmental and astrobiological implications

The indigenous microbiome of the Dhala area represents a naturally evolved genetic reservoir with considerable potential for environmental biotechnology. Functional metagenomic analyses revealed a diverse range of genes involved in heavy-metal resistance, uranium transformation, oxidative stress response, membrane transport, nutrient cycling, and biomineralization, indicating that these microorganisms possess multiple complementary mechanisms for surviving and transforming radionuclides in situ. Similar metabolic capabilities have been widely recognized in microorganisms from uranium-rich environments, where metal reduction, biosorption, biomineralization, and extracellular electron transfer contribute to the immobilization and detoxification of uranium and other toxic metals [13,12,45,46,14]. Since these microbial communities have evolved under prolonged natural selection within a radionuclide-enriched geological setting, they may provide more robust and environmentally compatible biotechnological resources than engineered microbial systems for the remediation of uranium-contaminated soils, groundwater, mine tailings, and radioactive waste repositories.

Additionally, the present study also establishes the Dhala structure as an exceptional natural laboratory linking impact geology, geomicrobiology, environmental biotechnology, and astrobiology. Integration of whole-genome metagenomics with geological and geochemical observations demonstrates how long-term impact-related geological processes have shaped the ecological organization, evolutionary adaptation, and functional potential of a naturally uranium-associated microbiome. These findings advance our understanding of radionuclide biogeochemistry and deep biosphere evolution while reinforcing the importance of terrestrial impact structures as analogues for potentially habitable subsurface environments on early Earth and other rocky planetary bodies, particularly Mars [2,3,4,6,40,44,47,48].

## Conclusion

The present investigation provides the first whole-genome metagenomics-based profiling of the microbiome associated with the Dhala impact structure, India. Results revealed a cross-domain microbiome (up to species level) well adapted to constant radiotoxicity. Exploration of this unique system revealed 58 genes involved in uranium biomineralization, tolerance, and cellular efflux. Findings from this study confirmed that the ancient impact structures on earth are exceptional evolutionary laboratories possessing untapped genomic landscapes for metal homeostasis. Beyond highlighting geosphere-biosphere interactions, the latest identified genomic repertoire offers a strong biotechnological blueprint. Exploiting these extremophilic pathways provides an eco-friendly, highly effective base for the development of next-generation microbial bioremediation strategies to bioremediate anthropogenic nuclear waste and uranium contamination at the global level.

Furthermore, the discovery of metabolically versatile cross-domain microbial communities possessing extensive stress-response systems, heavy-metal resistance, and biomineralization pathways strengthens the hypothesis that comparable impact-generated environments on early Mars and other rocky planetary bodies may have supported microbial ecosystems. Therefore, the taxonomic signatures, functional genes, and metabolic pathways identified in the Dhala samples provide valuable terrestrial analogues for interpreting potential biosignatures preserved within extraterrestrial impact-generated rocks and subsurface hydrothermal systems.

## Acknowledgements

The authors are thankful to Centyle Biotech Private Limited, India, for providing whole-genome metagenomics sequencing services. The authors are also thankful to the Department of Earth and Planetary Sciences, Nehru Science Centre, University of Allahabad, India, and the Department of Zoology, University of Allahabad, India.

## Author contributions

K., & P.K. conceptualization, investigation, resources, supervision, formal analysis, writing-original draft. S.D., & A.K.S investigation, writing-review & editing. J.K.P investigation, resources, supervision, writing-review & editing.

